# G9a/GLP methyltransferases inhibit autophagy by methylation-mediated ATG12 protein degradation

**DOI:** 10.1101/2021.02.05.430008

**Authors:** Chang-Hoon Kim, Kyung-Tae Park, Shin-Yeong Ju, Cheol-Ju Lee, Sang-Hun Lee

## Abstract

Previous studies have shown that G9a, a lysine methyltransferase, inhibits autophagy by repressing the transcription of autophagy genes. Here, we demonstrate a novel mechanism whereby G9a/GLP inhibit autophagy through post-translational modification of ATG12, a protein critical for the initiation of autophagosome formation. Under non-stress conditions, G9a/GLP directly methylate ATG12. The methylated ATG12 undergoes ubiquitin-mediated protein degradation, thereby inhibiting autophagy induction. By contrast, under stress conditions that elevate intracellular Ca^2+^ levels, the activated calpain system cleaves the G9a/GLP proteins, leading to G9a/GLP protein degradation. The reduced G9a/GLP levels allow ATG12 to accumulate and form the ATG12-ATG5 conjugate, thus expediting autophagy initiation. Collectively, our findings reveal a distinct signaling pathway that links cellular stress responses involving Ca^2+^/calpain to G9a/GLP-mediated autophagy regulation. Moreover, our model proposes that the methylation status of ATG12 is a molecular rheostat that controls autophagy induction.

## INTRODUCTION

Macroautophagy (hereinafter autophagy), which is evolutionarily conserved from yeast to mammals, is a key process for the turnover and recycling of long-lived proteins, cellular macromolecules, and damaged organelles ^1, 2^. Autophagy is initiated by the formation of double membrane structures called autophagosomes and mediated by autophagy-related (ATG) proteins. Defects in autophagy are associated with various disorders, such as cancer and neurodegenerative diseases ^3, 4^. Therefore, elucidating how autophagy is controlled is crucial for developing therapeutics for these disorders.

“Under non-stress and nutrition-rich conditions, autophagy induction is suppressed by mTOR that is activated by PI3K/AKT pathway. Therefore, mTOR is a central intracellular transducer that regulates autophagsome formation. However, under stress conditions that promote autophagy induction, mTOR is inhibited by AMPK and HIF that are activated by starvation and hypoxic condition, respectively. Recent studies have suggested that intracellular Ca^2+^ levels also play an important role in inducing autophagy via the CaMKK-β-AMPK-mTOR or AMPK-independent pathways ^7–9^. Calpains belong to the papain family of Ca^2+^-dependent cysteine proteases ^10^ and are required for autophagy in mammalian cells ^11, 12^. This raises the possibility that calpains might regulate autophagy stimulated by increased intracellular Ca^2+^. However, the molecular mechanisms by which calpains regulate autophagy associated with elevated intracellular Ca^2+^ level remain unknown.

Post-translational modifications (PTMs) play an important role in regulating protein stability, targeting specific subcellular compartments, protein–protein interactions, and other functionalities ^13^.In fact, the function of ATG proteins is modulated by PTMs such as phosphorylation, ubiquitination, and lipidation ^14^. However, there have been no reports that ATG proteins are methylated. Euchromatic histone lysine methyltransferase 2 (EHMT2/G9a) and G9a-like protein (GLP/EHMT1) were originally identified as lysine methyltransferases (KMT) that catalyze histone H3K9 methylation to generate repressive histone codes ^15^. G9a and GLP (hereafter, G9a/GLP) interact with their substrates through their SET domains and function primarily as a heteromeric complex *in vivo* ^16^. Intriguingly, previous studies have demonstrated that G9a also methylates non-histone proteins to control their stability and functions ^17–19^.

ATG proteins play a central role in regulating autophagy by controlling autophagosome formation. Upon autophagy induction, ATG12 is conjugated to ATG5, thereby having (or acquiring)?a ubiquitin-like E3 ligase activity in the cytoplasm ^20^. In parallel, a cytoplasmic microtubule-associated protein 1A/1B light chain 3 (LC3) is converted into light chain 3I (LC3I) by ATG4, which becomes subsequently lipidated by ATG3/7 ^21^. The lipidated form, LC3II, is incorporated into the autophagosome by the ATG12-ATG5-ATG16L1 complex and is generally used as a marker for autophagy ^22^.

Recent evidence indicates that epigenetic suppression of autophagy gene expression is an important regulatory mechanism of autophagy induction. The histone deacetylase inhibitors sodium butyrate and suberoylanilide hydroxamic acid (SAHA) induce autophagy and autophagic cell death ^23–25^. In line with these findings, histone methylation markers such as H3K9me2 and H3K27me3 have been shown to be an integral part of the autophagy process ^26, 27^. Moreover, inhibition of G9a/GLP with BIX01294 activates autophagy by increasing the transcription of numerous autophagy genes ^26, 28^. Despite the accumulating evidence, whether histone-modifying enzymes such as G9a/GLP and the JmjC domain-containing histone demethylase 2/lysine-specific demethylase 4 (JMJD2/KDM4) family regulate autophagy induction by directly modifying ATG proteins have not been explored.

In this study, we provide compelling evidence that under non-stress conditions G9a/GLP inhibit autophagy by directly methylating ATG12, which subsequently undergoes ubiquitination and degradation. However, under stress conditions, calpains activated by increased cytosolic Ca^2+^ levels cleave G9a/GLP, which results in the accumulation of ATG12, the ATG12-ATG5 conjugate, and LC3II and promotes autophagy induction. Our study therefore reveals a novel mechanism of autophagy regulation by G9a/GLP.

## RESULTS

### Cellular stress–induced autophagy reduces G9a/GLP protein levels via the Ca^2+^/calpain system

Cellular stresses induced by starvation or by inhibition of mTOR by rapamycin have been shown to stimulate autophagy through elevated Ca^2+^ signaling ^29^. Interestingly, inhibition of G9a methyltransferase activity has been reported to induce autophagy ^26, 28, 30–35^. Therefore, we looked into a causal link between Ca^2+^ signaling and G9a in autophagy induction. Endogenous G9a levels gradually declined in the presence of autophagy inducers such as the Ca^2+^ ionophore A23187, starvation by EBSS medium, and the mTOR inhibitor Torin1 (Fig. 1a). A reduction in G9a protein levels upon autophagy induction by Torin1 and starvation in HEPES ^36^ and EBSS media was further confirmed by immunofluorescence staining (Fig. 1b). Moreover, under starvation in HEPES medium, the reduction in G9a protein level was accompanied by increased LC3BII and decreased p62/SQSTM1 levels (Fig. 1c). Interestingly, the Ca^2+^ chelator BAPTA/AM significantly prevented the decrease in the G9a protein level (Fig. 1b,c), suggesting that Ca^2+^ signaling plays an important role in G9a reduction during autophagy. Similarly, the level of exogenously expressed GFP-G9a in 293T cells was also diminished by autophagy (Supplementary Fig. 1a), indicating that G9a downregulation occurs at the post-translational level. In support of our conclusion, the *G9a* mRNA level was not altered by EBSS or UNC0638 treatment (Supplementary Fig. 1b).

**Fig. 1.**
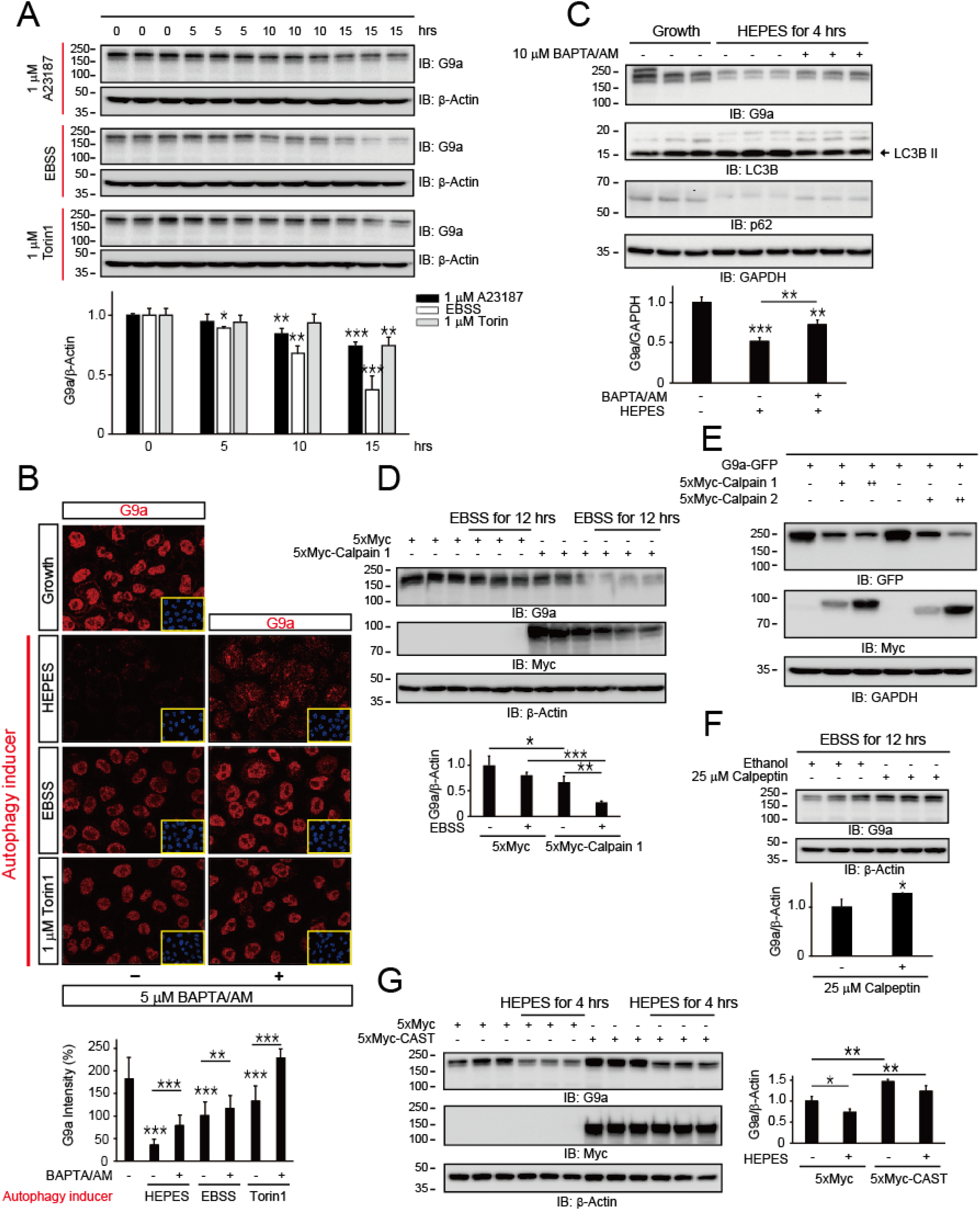
Autophagy downregulates G9a protein level through calpain activation. **a,** Autophagy induced by elevated cytoplasmic Ca^2+^ (1 µM A23167), nutrient starvation (EBSS), and metabolic stress (1 µM Torin1) decreased the G9a protein level. For starvation-induced autophagy, Hela cells were washed twice with PBS and incubated with EBSS for the indicated time. An mTOR inhibitor (Torin1) and a Ca^2+^ selective ionophore (A23187) were solubilized in dimethyl sulfoxide and diluted with DMEM containing 10% FBS to achieve the indicated final concentrations. **b,** Confocal immunofluorescence (IF) revealed that the endogenous G9a protein level was diminished during starvation in HEPES and EBSS media and metabolic stress (Torin1)-induced autophagy. However, sequestration of Ca^2+^ by 5 µM BAPTA/AM prevented the reduction in G9a protein in Hela cells. The intensity of the G9a protein was quantified using the mean fluorescence intensity (MFI) of individual cells (n = 3 [40–80 per group]). **c,** The G9a protein level was diminished during serum starvation (HEPES)-induced autophagy in Huh7 cells. However, in the presence of a Ca^2+^-chelating agent (10 µM BAPTA/AM), the G9a protein level was significantly elevated. **d,** Calpain 1 expression downregulated the endogenous G9a protein in 293T cells. **e,** Calpain 1 and 2 expression downregulated exogenous G9a-GFP in 293T cells. **f,** Inhibition of calpains by calpeptin, a cell permeable calpain inhibitor, elevated the endogenous G9a protein level in 293T cells under starvation in EBSS medium. **g,** Expression of CAST/Calpastatin, an endogenous calpain inhibitory protein, prevented a reduction in the G9a protein level in Hela cells under growth and starvation (HEPES) conditions. All experiments were repeated at least three times, and statistical analyses were performed using student’s t tests to compare groups (mean ± SEM of n = 3 replicates; *p < 0.05; **p < 0.01; ***p < 0.001).

Calpains are activated by Ca^2+^ and known to induce autophagy ^11, 12^. Indeed, the expression of calpain 1 significantly decreased endogenous G9a/GLP protein levels under nutrient-rich and starvation conditions (Fig. 1d; Supplementary Fig. 2a). We further confirmed the calpain-mediated downregulation of exogenously expressed G9a/GLP proteins (Fig. 1e; Supplementary Fig. 2b). Conversely, inhibition of calpains with calpeptin, a cell-permeable calpain inhibitor, increased endogenous G9a/GLP protein levels under starvation in EBSS medium (Fig. 1f; Supplementary Fig. 2c). Likewise, the expression of CAST/Calpastatin, an endogenous calpain inhibitory protein, upregulated G9a/GLP protein levels under non-stress and starvation (HEPES) conditions (Fig. 1g; Supplementary Fig. 2d). Taken together, our results indicate that the calpain system downregulates G9a/GLP protein levels upon autophagy induction.

### G9a cleavage by calpains leads to degradation of G9a through the ubiquitin-proteasome system (UPS)

Because G9a protein level was reduced by calpain expression, we examined whether G9a directly interacts with calpains and found that calpain 1 and 2 indeed associate with G9a in immunoprecipitation (IP) assays (Fig. 2a). To determine whether G9a is directly cleaved by calpains, we expressed G9a fused with a C-terminal tagged GFP in 293T cells and detected the C-terminus fragments of G9a protein in Western blots using a GFP antibody. Several cleaved G9a fragments appeared when the Ca^2+^ ionophore A23187 was added or calpains were expressed (Fig. 2b). These cleaved forms of G9a accumulated in the presence of MG132, a proteasome inhibitor (Fig. 2b,f), suggesting that G9a cleavage by calpains leads to G9a protein degradation via the UPS. As expected, calpain 1 and 2 expression augmented the ubiquitination level of G9a (Fig. 2c). Consistently, calpain activation using the Ca^2+^ ionophore A23187 increased G9a ubiquitination, whereas the Ca^2+^ chelator BAPTA/AM suppressed this effect (Fig. 2d). Ca^2+^, calpain 1 and 2, and MG132 produced similar effects on GLP cleavage patterns (Supplementary Fig. 2e). In line with the results, MG132 prevented the expected decrease in full-length and cleaved G9a levels under starvation in HEPES medium (Supplementary Fig. 3). Our findings collectively indicate that G9a/GLP proteins cleaved by calpains are destined for UPS-mediated protein degradation during autophagy.

**Fig. 2.**
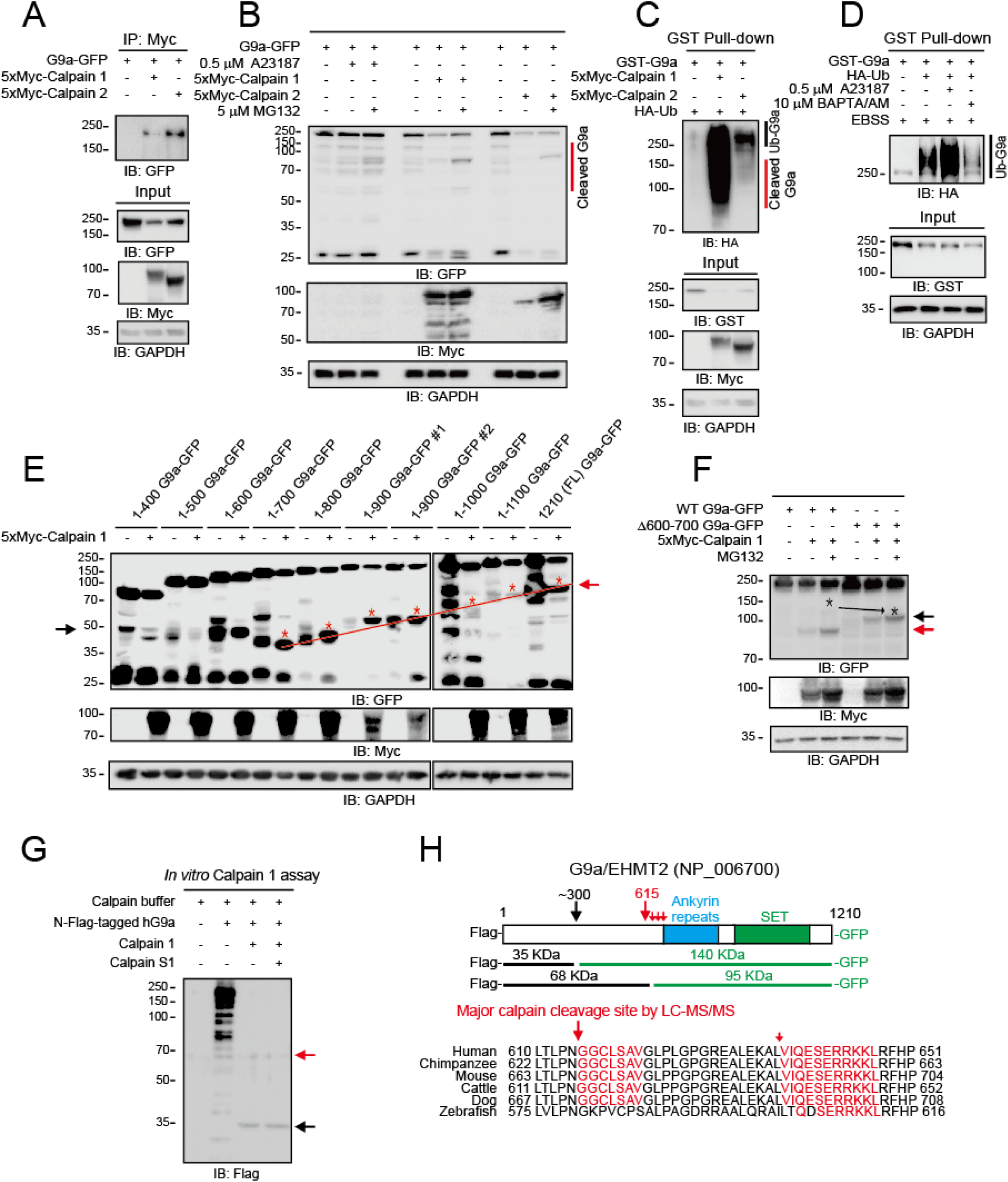
Calpains cleave G9a protein for ubiquitin-mediated degradation. **a,** Immunoprecipitation (IP) assays show that G9a bound to calpain 1 and 2. 293T cells were co-transfected with 5xMyc-calpain and G9a-GFP plasmids, and samples were prepared as described in the STAR Methods. To prevent G9a cleavage by calpains, the calpain inhibitor calpeptin (10 µM), protease inhibitors, and EGTA (1 mM) were added. **b,** G9a was cleaved by elevated Ca^2+^ influx (A23187) and the expression of calpains in 293T cells. The accumulation of cleaved G9a fragments by MG132 indicates that the cleaved G9a was degraded by the UPS. **c,** Expression of calpain 1 or calpain 2 promoted G9a ubiquitination in 293T cells. 293T cells were co-transfected with the indicated constructs. **d,** Elevated Ca^2+^ influx by A23187 promoted G9a ubiquitination, whereas Ca^2+^ chelation by BAPTA/AM reduced G9a ubiquitination under starvation in EBSS medium. GST pull-down assays were performed to monitor the ubiquitination level of G9a-GST after 12 hours of EBSS starvation. **e,** Calpain cleavage sites in G9a are located between aa 600 and aa 700. **f,** The Δ600–700 G9a-GFP mutant lacking the calpain cleavage sites was more resistant than WT G9a-GFP to calpain 1, as indicated by the lack of cleavage products (red arrow) and a higher level of intact G9a protein. The cleavage products of the Δ600–700 G9a (black arrow) suggest the presence of an additional calpain cleavage site at the N-terminus of G9a. **g,** An *in vitro* calpain cleavage assay confirmed that G9a has two cleavage sites. The main site is between aa 600 and aa 700. An additional N-terminal cleavage site is near aa 300. Recombinant Flag-tagged hG9a, calpain 1, and calpain S1 (a calpain 1 co-activator) were incubated for 10 min at 37°C, and the reaction products were subjected to Western blot analysis. **h,** A schematic diagram of the G9a cleavage sites and fragments in Fig. 2f, g and Supplementary Fig. 5. The calpain cleavage sequences in G9a are highly conserved across vertebrates.

### Determination of calpain cleavage sites in G9a

To confirm our findings above, we sought to identify the calpain cleavage site in G9a. The molecular weight of full-length G9a-GFP is around 200 kDa. A C-terminal G9a cleavage fragment (95 kDa) including the GFP protein was produced in the presence of calpains (Fig. 2b). Therefore, the molecular weight of the cleaved G9a fragment was estimated to be approximately 70 kDa when GFP (26 kDa) was excluded. Thus, the cleavage site in G9a could be located between amino acid (aa) 600 and aa 700. To test this idea, we generated several G9a truncation mutants with a C-terminal GFP tag. The G9a mutants produced cleaved fragments regardless of exogenous calpain 1 expression, possibly due to endogenous calpain activity (Fig. 2e). Nonetheless, when all the cleaved G9a fragments were connected by the line highlighted in red, it terminated at the 1-700 G9a-GFP mutant (Fig. 2e), indicating that the calpain cleavage site in G9a is between aa 600 and aa 700. This interpretation was supported by the observation that fluorescence intensity from the 1-700 G9a-GFP mutant to the full-length G9a-GFP construct was significantly diminished in the presence of calpain 1 expression (Supplementary Fig. 4). Based on the intensity of the cleaved bands of G9a truncation mutant proteins, the major cleavage sites for calpains in the G9a protein are located at aa 600–700.

We next performed LC-MS/MS analyses using C-terminal GST-tagged G9a to determine specific calpain cleavage sites. The main cleavage site in G9a starts with a GGCLSAV sequence (615 position, Fig. 2h; Supplementary Fig. 5). To test whether a G9a mutant deficient in the observed calpain cleavage sites between aa 600 and aa 700 is resistant to calpain-induced proteolysis, we produced a G9a mutant lacking aa 600 to 700 (Δ600–700 G9a) and found that the Δ600–700 G9a protein was more resistant to calpain 1-mediated cleavage than the wild type (WT) G9a, as indicated by low levels of the cleaved forms (Fig. 2f, red arrow). Fortuitously, throughout the course of our studies on calpain-induced Δ600–700 G9a proteolysis, we identified an additional cleavage site for calpains in G9a at aa ∼300. The N-terminal cleavage of Δ600–700 G9a-GFP compared with WT G9a-GFP is clearly visible in Fig. 2f (black arrow). We further confirmed the existence of N-terminal cleavage site from 1–400 G9a-GFP mutant to 1–600 G9a-GFP mutant (Fig. 2e, black arrow on left side). The N-terminal cleavage in G9a by calpain 1 was negligible in the presence of the major calpain cleavage sites in WT G9a. The N-terminal cleavage becomes apparent only when major calpain cleavage sites are absent in Δ600–700 G9a (Fig. 2f, black asterisk and arrow). However, N-terminal cleavage is not required for G9a protein degradation, as shown by the lack of response to calpain 1 (Supplementary Fig. 4). An *in vitro* calpain cleavage assay using purified proteins confirmed the two cleavage sites for calpains in the G9a protein (Fig. 2g,h, red and black arrows). Taken together, our data indicate that the calpain cleavage sites in aa 600–700 are important for G9a protein degradation even though G9a has an additional N-terminal cleavage site for calpains.

### G9a/GLP inhibit autophagy induction

Given our finding that autophagy induction involves G9a downregulation, we hypothesized that G9a might play an inhibitory role in autophagy induction. Therefore, we sought to determine the function of G9a in autophagy. Intriguingly, both WT G9a and the Δ600–700 G9a mutant decreased the levels of the ATG12-ATG5 conjugate and LC3BII while increasing the p62/SQSTM1 level under starvation conditions (Supplementary Fig. 6a). Likewise, GLP expression reduced the ATG12-ATG5 conjugate level (Supplementary Fig. 6b), suggesting that G9a/GLP proteins indeed inhibit autophagy induction. Conversely, calpain 1 and calpain 2, which induce the UPS-mediated G9a/GLP protein degradation (Fig. 2; Supplementary Fig. 2), elevated the formation of the ATG12-ATG5 conjugate and LC3BII under starvation in HEPES medium (Supplementary Fig. 6c). To determine the role of G9a on ATG12-ATG5 conjugate formation, we used ATG5-null cells. In the absence of ATG5 expression, inhibiting G9a with UNC0638 had no effect on LC3BII formation, whereas the introduction of ATG5 recovered UNC0638-mediated LC3BII formation in ATG5 −/− cells (Supplementary Fig. 6d), suggesting that G9a regulates autophagosome formation by acting on the ATG12-ATG5 conjugation formation step or in its downstream pathway.

### G9a/GLP methylates ATG12 in the cytoplasm

G9a has been thought to locate primarily in the nucleus. However, accumulating evidence indicates that the G9a protein localizes in both the nucleus and the cytoplasm ^37–39^. A recent study in particular revealed that the localization of endogenous G9a to the cytoplasmic compartment requires the exclusion of exon 10 (E10) from the *G9a* mRNA ^38^. Consistent with this study, we were able to detect cytoplasmic G9a protein by immunofluorescence staining (Supplementary Fig. 7a). Moreover, we observed alterations in the G9a localization patterns by using Leptomycin B, a highly specific nuclear export inhibitor ^40, 41^. Apparently, Leptomycin B induced the nuclear accumulation of G9a while causing disappearance of cytoplasmic G9a (Supplementary Fig. 7b,c), indicating that G9a shuttles between the nucleus and the cytoplasm.

These findings suggest that G9a is cleaved by calpains in the cytoplasm (Fig. 1, 2; Supplementary Fig. 4), and that G9a may play an essential role in preventing autophagy under non-stress conditions by methylating key ATG proteins such as ATG12 and ATG5. Therefore, we investigated whether G9a directly methylates ATG12 and ATG5. Our GST-pulldown assays revealed that G9a interacts with ATG12, ATG5, and LC3B (Fig. 3a). In addition, IP assays revealed that endogenous G9a associates with endogenous ATG12 and ATG5 (Fig. 3b). Interestingly, JMJD2C, which demethylates H3K9me3 and H3K36me3 ^42–44^ and therefore counteracts G9a activity ^45, 46^, also interacted with ATG12, ATG5, and LC3B (Supplementary Fig. 8a). This suggests that the JMJD2 family of proteins might promote autophagy by demethylating the ATG proteins. In an *in vitro* methyltransferase assay using purified GST-fusion proteins, the enzymatically active G9a SET protein (G9a-SET) methylated only ATG12, not ATG5, LC3B, or ATG3 (Supplementary Fig. 8b, see *). The same observation was made with GLP (Supplementary Fig. 8c, see *). These results support the idea that G9a methylates both histone and non-histone proteins ^17^.

**Fig. 3.**
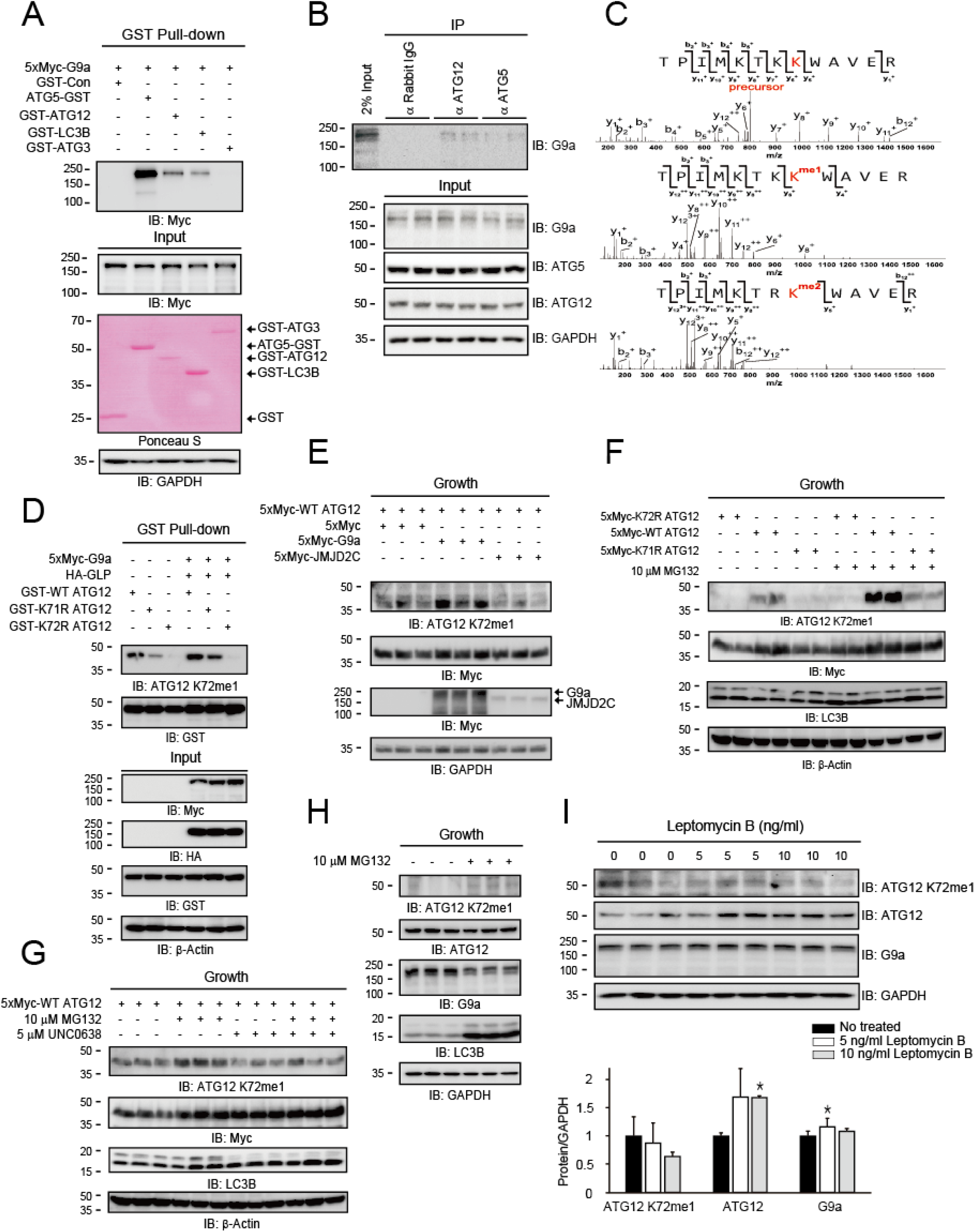
G9a methylates K72 residue in ATG12. **a,** GST pull-down assays show a direct interaction of G9a with ATG12, ATG5, and LC3B. **b,** IP assays reveal a specific interaction of endogenous G9a with ATG12 and ATG5 in 293T cells. **c,** Mass spectrometry analysis confirmed mono- and di-methylation at K72 residue in ATG12. The GST-ATG12-B used in Fig. 3d was subjected to Glu-C endoproteinase digestion and LC-MS/MS analysis. **d,** K72R ATG12 was not methylated by either endogenous or exogenous G9a/GLP, suggesting that G9a and GLP specifically methylate K72 in ATG12. WT ATG12 was more extensively mono-methylated by G9a/GLP than K71R ATG12 in 293T cells. **e,** K72 mono-methylation (K72me1) by endogenous G9a/GLP was reversed by JMJD2C expression in Hela cells. **f,** MG132 caused the accumulation of ATG12 K72me1 in WT ATG12 and K71R ATG12 in Hela cells, confirming that K72 methylation led to ATG12 degradation by the UPS pathway. **g,** Inhibition of G9a/GLP activity by UNC0638 reduced the ATG12 K72me1 level in Hela cells. **h,** MG132 increased the ATG12 K72me1 level in Hela cells under non-stress conditions. **i,** Leptomycin B decreased the ATG12 K72me1 level but increased the protein levels of the ATG12-ATG5 conjugate in the cytoplasm and G9a in the nucleus in Hela cells (mean ± SEM of n = 3 replicates; *p < 0.05).

### K72 residue in ATG12 is methylated by G9a

Human ATG12 has 12 lysine (K) residues that could serve as potential G9a methylation sites. To identify the ATG12 residue(s) methylated by G9a, we generated four GST-ATG12 fusion proteins containing distinct K residues (Supplementary Fig. 8d, upper panel). An *in vitro* G9a methyltransferase assay revealed that a GST-ATG12 fragment protein containing the _63_GDTPIMKTKKWAVER_77_ peptide (fragment B) was specifically methylated by G9a (Supplementary Fig. 8d, see *). Next, to determine which K residue in the _63_GDTPIMKTKKWAVER_77_ peptide was methylated, each of the K residues in the peptide was sequentially replaced with arginine (R) (Supplementary Fig. 8e, upper panel). The methylated band detected by ^3^H-SAM autoradiography disappeared by a substitution at K72 (K72R) but not by K69R or K71R (Supplementary Fig. 8e), indicating that K72 is a specific residue methylated by G9a. Interestingly, a K71R mutation yielded a very intense methylation band (Supplementary Fig. 8e, see **), indicating that R_71_K_72_ is a more favorable target motif for G9a methyltransferase than the WT motif, K_71_K_72_. These findings were further confirmed in full-length ATG12 (Supplementary Fig. 8f, see *). Consistent with Supplementary Fig. 8e, methylation was enhanced by K71R substitution (Supplementary Fig. 8f, see **). Furthermore, when an additional K89R substitution was introduced in the presence of the K71R mutation, two favorable RK motifs (R_71_K_72_ and R_89_K_90_) were produced in ATG12 protein, and ATG12 methylation by G9a was further increased (Supplementary Fig. 8f, see ***). Notably, no methylation was detected with the ATG12 proteins containing the K72R mutation (Supplementary Fig. 8f, lanes 4–6).Consistently, LC-MS/MS analyses of WT GST-ATG12-B revealed that K72 is not methylated in the absence of the G9a-SET enzyme (Fig. 3c, upper panel). However, K72 was mono-methylated in _65_TPIMKT**K_71_K_72_**WAVER_77_ (Fig. 3c, middle panel) and di-methylated in _65_TPIMKT**R_71_K_72_**WAVER_77_ (Fig. 3c, bottom panel) in the presence of the G9a-SET enzyme, supporting the idea that the R_71_K_72_ motif is a more favorable target for G9a. Collectively, our results indicate that K72 in ATG12 is a specific G9a methylation site.

### K72me1 in ATG12 is coupled to ATG12 protein degradation via the UPS pathway under non-stress conditions

Since we could not detect endogenous K72 methylation using mass spectrometry, possibly due to a high turnover rate of methylated ATG12 in cells, we decided to examine K72 methylation in ATG12 inside cells. Therefore, we raised specific antibodies against a peptide in which K72 is mono-methylated (K72me1: IMKTKK(me)WAVER, Supplementary Fig. 9). Among 4 batches, antibody batches 3 and 4 (K72me1 #3 and #4) exhibited high specificity and sensitivity (Supplementary Fig. 9). As expected, K72me1 was not detected in K72R ATG12 even when G9a was overexpressed, and K72me1 was clearly detected in WT ATG12 and K71R ATG12 even in the absence of G9a/GLP overexpression (Fig. 3d). However, to our surprise, WT ATG12 was more mono-methylated than K71R ATG12 (Fig. 3d, f), possibly due to a rapid transition of K72me1 to K72me2 inside cells, thereby depleting K72me1inside cells. As shown above, the K71R ATG12 mutant was mostly di-methylated (Fig. 3c) Consistently, G9a/GLP expression increased the K72me1 level (Fig. 3d,e), whereas G9a/GLP inhibition through UNC0638 diminished it (Fig. 3g). Notably, K72me1 produced by endogenous G9a/GLP was removed by JMJD2C (Fig. 3e), further verifying the specificity of the antibody. MG132 upregulated the K72me1 level under non-stress conditions (Fig. 3f–h), indicating that ATG12 containing K72me1 undergoes protein degradation by the UPS. Moreover, in agreement with the idea that the cytoplasmic G9a methylates ATG12 (Supplementary Fig. 7), Leptomycin B significantly decreased the ATG12 K72me1 level while increasing the ATG12-ATG5 conjugate and G9a levels (Fig. 3i). Thus, these findings indicate that ATG12 in the cytoplasm was stabilized due to a lack of methylation by G9a, and that G9a in the nucleus was protected from the cytoplasmic calpains.

### Reduced levels of G9a/GLP increasethe ATG12-ATG5 conjugate level

We next examined the relationship between G9a/GLP and the ATG12-ATG5 conjugate levels during autophagy. When autophagosomes fuse with lysosomes to form autolysosomes, ATG12 isdownregulated through lysosomal degradation of the ATG12-ATG5-ATG16L1 complex. As G9a/GLP levels declined, the ATG12-ATG5 conjugate level was elevated at the early stages of autophagy, but was later decreased by lysosomal degradation of the ATG12-ATG5 conjugate (Fig. 4a–c). Interestingly, the GLP protein was transiently upregulated under starvation in EBSS medium (Fig. 4c), possibly due to a negative feedback mechanism involving the upregulation of *GLP* mRNA expression (Supplementary Fig. 1b). The level of endogenous ATG12 K72me1, which is produced by G9a, exhibited strikingly similar patterns to those of endogenous G9a and GLP (Fig. 4a–c). This observation suggests that G9a/GLP downregulate the ATG12-ATG5 conjugate level by methylating K72 in ATG12.

**Fig. 4.**
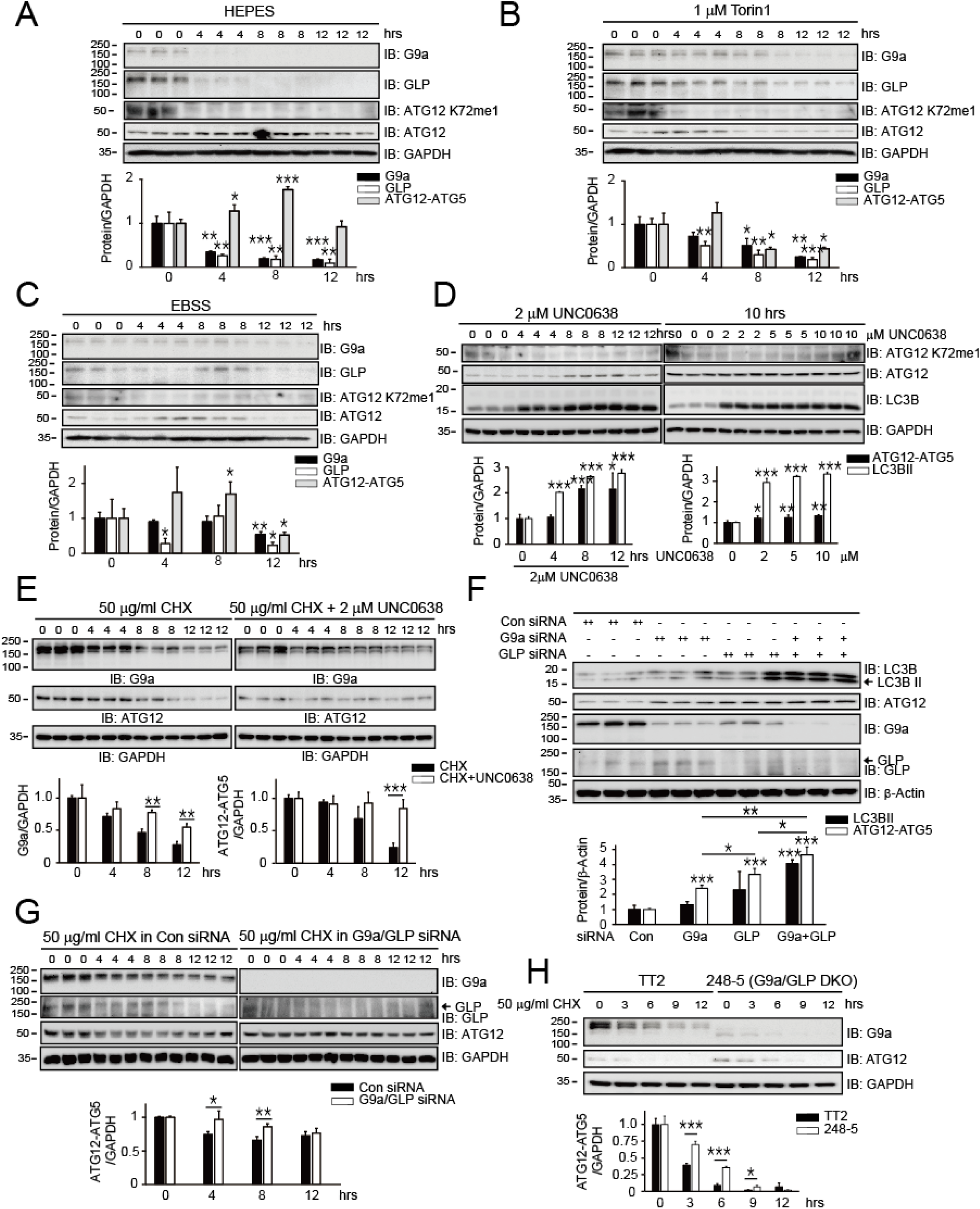
G9a/GLP downregulate the ATG12-ATG5 conjugate level. **a,** During serum starvation (HEPES)-induced autophagy in Hela cells, the levels of both the G9a and GLP proteins declined, whereas the ATG12-ATG5 conjugate level was transiently elevated. **b,** Autophagy induced by mTOR inhibition led to a reduction in the G9a and GLP protein levels, with a concomitant increase in the ATG12-ATG5 level in Hela cells. **c,** Although the GLP protein was transiently upregulated, EBSS gradually decreased G9a/GLP protein levels. The ATG12-ATG5 level was steadily augmented for 8 hours in Hela cells. **d,** G9a inhibition with UNC0638 elevated the ATG12-ATG5 conjugate level and promoted the transformation of LC3BI to LC3BII in a time- and dose-dependent manner in Hela cells. **e,** Inhibition of G9a (2 µM UNC0638) in the presence of 50 µg/ml of CHX stabilized the ATG12-ATG5 conjugate compared with the control treated with only CHX in Hela cells. **f,** Silencing G9a and/or GLP expression additively increased the formation of the ATG12-ATG5 conjugate and the conversion of L3CBI to LC3BII in U2OS cells. **g,** Silencing both G9a and GLP expression significantly stabilized the ATG12-ATG5 conjugate in U2OS cells. **h,** The ATG12-ATG5 conjugate had an extended half-life in G9a/GLP double-knockout embryonic stem cells (248-5 cell line) compared with the WT TT2 cell line. All experiments were repeated at least three times, and statistical analyses were performed using student’s t tests to compare groups (mean ± SEM of n = 3 replicates; *p < 0.05; **p < 0.01; ***p < 0.001).

To test this idea, we inhibited G9a with UNC0638.The inhibition of G9a elevated the formation of the ATG12-ATG5 conjugate and LC3BII in a time-dependent fashion (Fig. 4d, left panel). Increasing the UNC0638 concentration further augmented the formation of the ATG12-ATG5 conjugate and LC3BII (Fig. 4d, right panel). When G9a was not inhibited, the ATG12 K72me1 level was significantly higher. However, at later time points with 2 µM UNC0638 or when high concentrations of the inhibitor were used for 10 hours, the ATG12 K72me1 level was increased (Fig. 4d), possibly due to the accumulation of the ATG12-ATG5 conjugate and the ATG12 K72me1 not being converted to the di-methylated ATG12 form (ATG12 K72me2, Fig. 3c). When cycloheximide (CHX) was added in conjunction with UNC0638, the level of the ATG12-ATG5 conjugate was stabilized (Fig. 4e), confirming that ATG12 methylation by G9a results in the conjugate turnover.

Because UNC0638 inhibits both G9a and GLP ^47^, we used an RNAi-based knockdown strategy to determine the contribution of each protein to autophagy. The expression of the G9a and GLP proteins was efficiently downregulated by the respective siRNA (Fig. 4f). The GLP siRNA displayed cross reactivity in that it also downregulated G9a expression (Fig. 4f, compare lanes 1–3 with lanes 7–9 in IB: G9a). Consistent with Fig. 4d, the G9a and GLP siRNA significantly increased autophagosome formation estimated by the levels of ATG12-ATG5 and LC3BII (Fig. 4f) and the ATG12 K72me1 level (Fig. 5e). In addition, silencing both G9a and GLP in the presence of CHX stabilized the ATG12-ATG5 conjugate like UNC0638 (Fig. 4g). Consistent with this, the ATG12-ATG5 conjugate in G9a/GLP double-knockout embryonic stem cells (248-5) exhibited an extended half-life compared with the control (TT2) (Fig. 4h). Taken together, both G9a and GLP contribute additively to the downregulation of the ATG12-ATG5 conjugate.

**Fig. 5.**
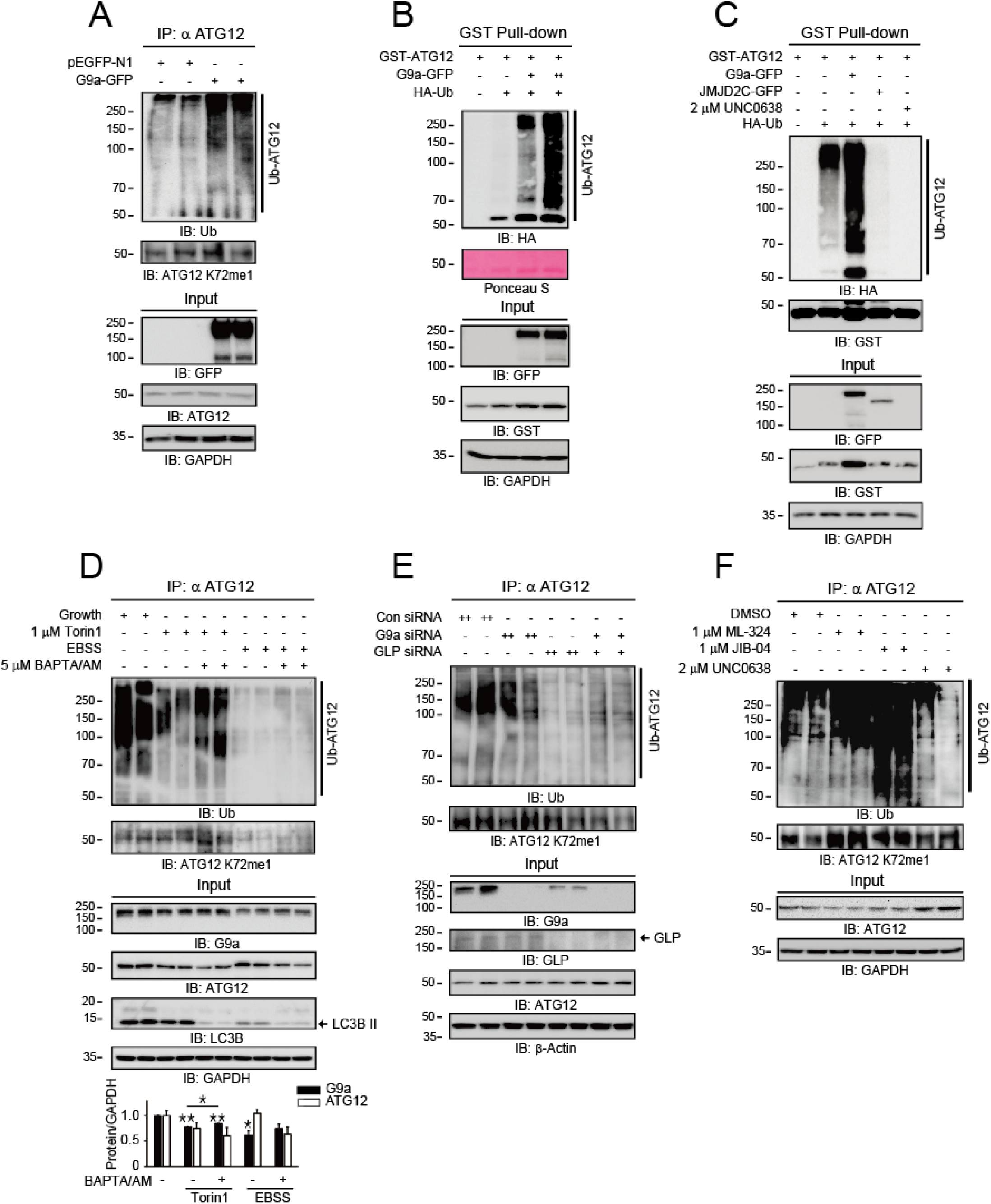
ATG12 methylated by G9a undergoes ubiquitination. **a,** G9a expression promoted endogenous ATG12 ubiquitination in Hela cells. Because the rabbit IgG heavy chains of the anti-ATG12 antibody overlap with the ATG12-ATG5 conjugate, an anti-rabbit IgG secondary antibody-conjugated HRP specific for light chains was used. **b,** GST pull-down assays show that the degree of ATG12 ubiquitination depended on the G9a expression level in 293T cells. **c,** G9a inhibition (2 µM UNC0638) and JMJD2C expression prevented ATG12 ubiquitination by counteracting the G9a/GLP activities in 293T cells. **d,** Autophagy induced by mTOR inhibition (1 µM Torin1) and starvation (EBSS) for 12 hours decreased the ubiquitination of endogenous ATG12. However, BAPTA/AM reversed this effect in Hela cells (mean ± SEM of n = 3 replicates; *p < 0.05; **p < 0.01). **e,** Silencing G9a and/or GLP expression significantly diminished endogenous ATG12 ubiquitination in Hela cells. **f,** G9a inhibition (2 µM UNC0638) suppressed endogenous ATG12 ubiquitination, whereas the inhibition of JMJD2 activity (1 µM ML-324 and 1 µM JIB-04) promoted endogenous ATG12 ubiquitination in Hela cells.

### K72 methylation in ATG12 by G9a is coupled to ATG12 ubiquitination

Consistent with our finding that K72me1 in ATG12 leads to ATG12 degradation via the UPS, G9a/GLP downregulate the ATG12-ATG5 conjugate level (Supplementary Fig. 6a,b). However, MG132 did not elevate the ATG12-ATG5 conjugate level (Supplementary Fig. 10). This could be due to degradation of the ATG12-ATG5 conjugate via a lysosome-mediated protein degradation pathway activated by MG132-induced autophagy ^48, 49^. In support of this possibility, MG132 increased the LC3BII level (Fig. 3f–h; Supplementary Fig. 10). Nonetheless, under non-stress conditions the endogenous ATG12 K72me1 level was clearly increased by MG132 (Fig. 3h), indicating that K72me1 in ATG12 serves as a degron for the UPS.

As anticipated, the ubiquitination of endogenous ATG12 protein was increased upon G9a expression (Fig. 5a), and the G9a-mediated ATG12 ubiquitination was further confirmed using exogenously expressed GST-ATG12 (Fig. 5b,c). As expected, both JMJD2C expression and G9a inhibition with UNC0638 inhibited ATG12 ubiquitination (Fig. 5c, lane 4 and 5). Autophagy induction with Torin1 and EBSS decreased the levels of endogenous ATG12 ubiquitination and G9a (Fig. 5d). On the contrary, BAPTA/AM, which modestly increases the G9a protein level under Torin1 and EBSS conditions (Fig. 1b), enhanced ATG12 ubiquitination along with G9a level (Fig. 5d). Furthermore, knockdown of G9a and/or GLP expression clearly reduced endogenous ATG12 ubiquitination, along with the ATG12 K72me1 level (Fig. 5e). In line with these observations, G9a inhibition with UNC0638 decreased ATG12 ubiquitination (Fig. 5f, compare lanes 1–2 with lanes 7–8). Although it was reported that very low level of the ATG12-ATG5 conjugate level is sufficient to induce autophagy in mouse fibroblast cells ^50^, our results in Fig. 4, 5 strongly suggest that G9a/GLP function as a rheostat that produce a significant effect on autophagy induction. Overall, our data suggest that ATG12 methylation by G9a/GLP leads to UPS-mediated ATG12 degradation.

### K72 residue is a methyl degron for ATG12 protein degradation

Our analyses of the methylation patterns of full-length ATG12 mutants (Supplementary Fig. 8f) indicated that the degree of methylation is in the order of K71R/K89R ATG12 > K71R ATG12 > WT ATG12 > K72R ATG12 > K72R/K89R ATG12 > K72R/K90R ATG12. Therefore, we further investigated how the degree of ATG12 methylation correlates with ATG12 protein degradation. In the absence of G9a expression, the ubiquitination levels of all the mutants were modest. However, ubiquitination of the ATG12 mutants showed an increasing pattern in accordance with their methylation propensity when G9a was expressed (Supplementary Fig. 11a, b), corroborating our finding that ATG12 methylation leads to its degradation by the UPS. Likewise, the degree of ATG5 ubiquitination was also proportional to the methylation propensity of the ATG12 mutants (Supplementary Fig. 11c), confirming that ubiquitination of the ATG5 protein is dictated by ATG12 methylation that prevents the formation of the ATG12-ATG5 conjugate. Therefore, the methylation status of K72 residue in ATG12 determines ATG12 and ATG5 protein degradation.

### The JMJD2 family promotes autophagy by counteracting G9a/GLP activity

Consistent with the results demonstrating a direct interaction between JMJD2C and ATG12 (Supplementary Fig. 8a) and the reduction of ATG12 K72me1 level by JMJD2C (Fig. 3e), JMJD2C expression augmented LC3BII and ATG12-ATG5 levels under autophagic conditions (Fig. 6a), indicating that JMJD2C antagonizes G9a activity to promote autophagy. Silencing JMJD2C produced a marginal reduction effect on the LC3BII level due to alow efficiency of JMJD2C knockdown (Fig. 6b) in Hela cells, which express much more JMJD2A and JMJD2B than JMJD2C according to the GEO Dataset (GSE108495, Fig. 6c). To overcome the limitation of siRNA, we used the JMJD2 inhibitors JIB-04 and ML-324 to inhibit the activity of the whole JMJD2 family (Fig. 6d). Inhibition of JMJD2 activity decreased the LC3BII level (Fig. 6e) and increased ATG12 ubiquitination and the ATG12 K72me1 level (Fig. 5f, lanes 3–6). Similarly, loss of JMJD2C increased ATG12 ubiquitination despite amodest JMJD2C knockdown (Fig. 6f). Furthermore, JMJD2C expression increased ATG12 protein stability (Fig. 6g), whereas ML-324, a JMJD2 family inhibitor, decreased the stability of the ATG12-ATG5 conjugate (Fig. 6h).

**Fig. 6.**
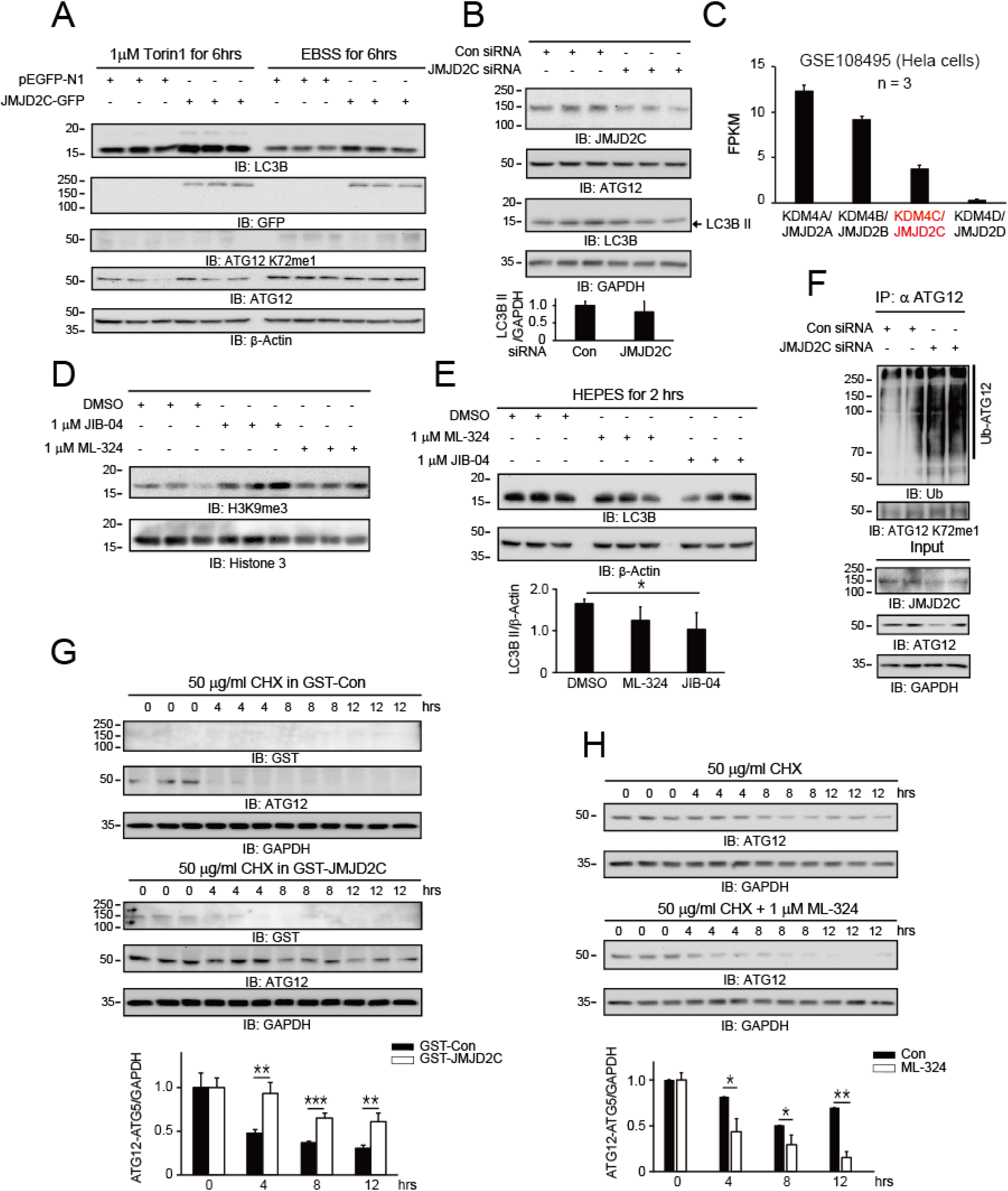
The JMJD2 family counteracts G9a/GLP activities on ATG12. **a,** JMJD2C expression in Hela cells promoted LC3BII conversion and increased the ATG12-ATG5 conjugate level during autophagy induced by mTOR inhibition (1 µM Torin1) and starvation (EBSS). **b,** Silencing JMJD2C expression in Hela cells modestly reduced both LC3BII and ATG12-ATG5 conjugate levels, indicating the active role of JMJD2C in autophagy induction. **c,** Gene expression analysis using GEO dataset GSE108495 revealed that the expression of JMJD2C was much lower than the expression of JMJD2A and JMJD2B in Hela cells. **d,** The JMJD2 family inhibitors JIB-04 and ML-324 blocked the activity of the histone demethylase JMJD2, thereby increasing the H3K9me3 level in Hela cells. **e,** The JMJD2 family inhibitors ML-324 and JIB-04 diminished LC3BI to LC3BII conversion during autophagy induced by HEPES medium in Hela cells, indicating that the JMJD2 family plays a role in promoting autophagy. **f,** Silencing JMJD2C expression increased endogenous ATG12 ubiquitination in Hela cells. **g,** JMJD2C expression stabilized the ATG12-ATG5 conjugate level in Hela cells. **h,** A JMJD2 family inhibitor (ML-324) diminished the stability of the ATG12-ATG5 conjugate in Hela cells. All experiments were repeated at least three times, and statistical analyses were performed using student’s t tests to compare groups (mean ± SEM of n = 3 replicates; *p < 0.05; **p < 0.01; ***p < 0.001). NS no significant difference (Student’s *t*-test).

Consistent with the previous observation that G9a overexpression reduced BIX01294-mediated LC3BII formation ^28^, JMJD2C expression produced a synergistic effect with UNC0638 in the formation of LC3BII in Hela cells (Supplementary Fig. 12a). Conversely, JMJD2C knockdown did not decreaseLC3BII formation, but silencing both G9a and GLP induced a much higher conversion of LC3BI to LC3BII (Supplementary Fig. 12b), supporting the idea that enhanced LC3BII formation results from the elevation of the ATG12-ATG5 conjugate level. Overall, our results clearly show that the JMJD2 family of proteins counteract G9a/GLP activity to promote autophagy.

### G9a promotes apoptosis through K72 methylation in ATG12 (or

We wanted to investigate the physiological significance of ATG12 methylation and regulation of autophagy induction by G9a.To this end, we reconstituted WT ATG12, a methylation-resistant K72R ATG12 mutant, and a hyper-methylation K71R ATG12 mutant in ATG12-deficient mouse embryonic fibroblast cells (ATG12−/− MEFs). As indicated by the LC3IIB level, basal autophagy was significantly elevated in cells expressing the methylation-resistant K72R ATG12 compared with those expressing WT ATG12 or K71R ATG12 (Fig. 7a). When autophagy was induced by Torin1, the hyper-methylated K71R ATG12 was the least effective in inducing autophagy, as shown by the LC3BII level (Fig. 7b), highlighting the importance of K72 methylation in ATG12 in preventing autophagy induction under stress conditions. Next, we wanted to examine physiological significance of K72methylation in ATG12Since autophagy is closely linked to apoptosis mostly preventing cell death program, we investigated how K72 methylation in ATG12 is related to apoptosis. we(ATG12−/− MEFs were detached under serum starvation conditions in EBSS and HEPES media. In this regard, previous studies have shown that G9a inhibition leads to apoptosis via increased autophagy ^28, 31, 35, 51^. When G9a and GLP activity was inhibited by 10 µM UNC0368 for 48 hours, ATG12-null MEF cells expressing methylation-resistant K72R ATG12 exhibited significantly lower levels of apoptosis than the cells expressing either WT ATG12 or the methylation-prone K71R ATG12, as characterized by cleaved p70 S6 kinase, an endogenous substrate of caspase 3 ^52, 53^ (Fig. 7c). These results clearly suggest that autophagy induction is critical for the survival of cancer cells that are resistant to apoptosis.

**Fig. 7.**
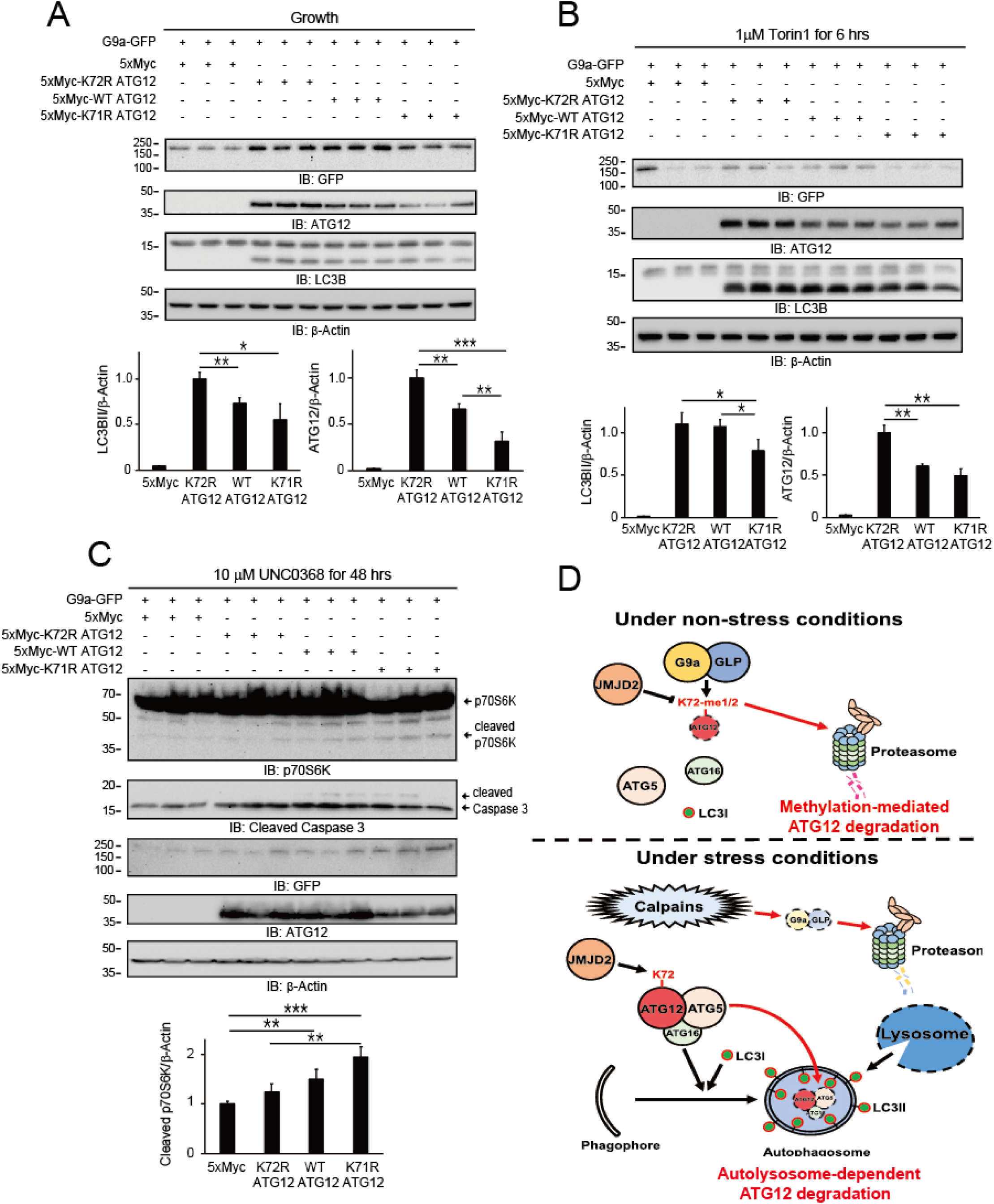
G9a regulates autophagy and apoptosis through K72 methylation in ATG12 **a,** Reconstitution of a methylation-resistant mutant of ATG12 (K72R) in ATG12-deficient MEFs promoted basal autophagy, whereas a hyper-methylation mutant of ATG12 (K71R) was less effective than WT ATG12 in inducing basal autophagy. **b,** Reconstitution of K72R ATG12 mutant in ATG12 −/− MEFs promoted autophagy when mTOR was inhibited (1 µM Torin1). **c,** Reconstitution of the methylation-resistant K72R ATG12 in ATG12-deficient MEFs conferred resistance to 10 µM UNC0368-induced apoptosis, whereas the expression of the hyper-methylated K71R ATG12 mutant in ATG12-deficient MEFs produced the opposite effect. **d,** Our model proposes that G9a/GLP inhibit autophagy by directly methylating ATG12. The methylated ATG12 subsequently undergoes methylation-mediated protein degradation under non-stress conditions. Under stress conditions, the Ca^2+^-activated calpain system downregulates the G9a/GLP proteins, thereby allowing the accumulation of ATG12, ATG12-ATG5 conjugate, and LC3II. Subsequent activation of lysosomal activity removes the ATG12-ATG5-ATG16L1 complex at a later stage of autophagy. All experiments were repeated at least three times, and statistical analyses were performed using student’s t tests to compare groups (mean ± SEM of n = 3 replicates; *p < 0.05; **p < 0.01; ***p < 0.001).

## DISCUSSION

In this study, we demonstrated that degradation of G9a/GLP by Ca^2+^-activated calpains is required for autophagy induction under stress conditions On the other hand, under non-stress conditions, G9a/GLP prevent autophagy induction by methylating ATG12, which reduces the level of the ATG12-ATG5 conjugate (Fig. 7d). Previous studies have shown that G9a prevents autophagy induction through the epigenetic transcriptional repression of autophagy genes ^26, 28^. Therefore, G9a/GLP methyltransferases prevent autophagy induction via dual mechanisms involving the transcriptional repression of autophagy genes and the post-translational modification of the ATG12 protein.

### The calpain system promotes G9a/GLP protein degradation

G9a and calpains display distinct subcellular localization patterns (Supplementary Fig. 4). G9a is known to localize in both the cytoplasm and the nucleus ^38^, whereas calpains are solely cytoplasmic proteins ^54–56^. Based on our studies (Fig. 1, 2; Supplementary Fig. 2, 4), the effects of calpain 1 and 2 on G9a/GLP protein degradation appear to be direct, strongly suggesting that calpains selectively cleave G9a/GLP localized to the cytoplasm. In agreement with the observations that the expression of calpain 1 downregulates G9a (Fig. 1d), and that the expression of CAST/Calpastatin, a calpain inhibitory protein, upregulates G9a (Fig. 1g), Leptomycin B, a highly selective of nuclear export, increases nuclear G9a (Supplementary Fig. 7b,c), indicating that G9a dynamically localizes in both the nucleus and the cytoplasm.In line with this Leptomycin B decreased the ATG12 K72me1 level, while increasing the ATG12-ATG5 conjugate level (Fig. 3i) by preventing the cytoplasmic localization of G9a (Supplementary Fig. 7b,c).

In accordance with previous studies that the calpain system promotes autophagy ^11, 12^, our findings show that calpain expression increases autophagy (Supplementary Fig. 6c). Nevertheless, there were conflicting reports that calpains inhibit autophagy by cleaving the ATG5 protein ^57, 58^. Therefore, it is conceivable that calpains might regulate the rate of autophagy by controlling the ATG5 level depending on the cellular context. This warrants further investigation. Alternatively, increased Ca^2+^ levels might activate yet unidentified pathways that degrade G9a/GLP. It is possible that the rate of G9a nuclear export to the cytoplasm is elevated under stress conditions. Despite the caveats, our results from Ca^2+^ chelation clearly indicate that Ca^2+^ plays a critical role in the induction of autophagy and activates the calpain system to degrade G9a/GLP and promote autophagy.

In addition, accumulating evidence indicates that the JMJD2 family of proteins also localizes in both the cytoplasm and the nucleus to a varying degree ^59–63^. In line with the reports, as shown in Fig. 3, 5, 6 and Supplementary Fig. 12, we found that JMJD2 family members demethylated ATG12 K72me1, thereby countering the methyltransferase activity of G9a on ATG12. G9a and JMJD2 proteins in the cytoplasm might compete constantly for ATG12 and the balance is tipped in favor of one or the other depending on cellular cues. Taken together, our results indicate that G9a dynamically shuttles between the nucleus and the cytoplasm, and that the cytoplasmic pool of G9a methylates ATG12 under non-stress conditions.

G9a protein is cleaved and degraded by calpains in response to elevated cytoplasmic Ca^2+^ levels, as evidenced by the effects of A23187 and BAPTA/AM on G9a ubiquitination (Fig. 2d). This result agrees well with a previous report that calpain-mediated protein cleavage is linked to proteasome-dependent protein degradation ^64^. This might explain the rapid degradation of G9a by the UPS. In this study, although we have not determined the identity of an E3 ubiquitin ligase that mediates the degradation of cleaved G9a protein, we speculate that it could be APC/C^Cdh1^, which is known to activate proteasomal degradation of G9a and GLP in response to the DNA damage response ^65^ and plays an important role in regulating the autophagic process ^66^.

### ATG12 methylation by G9a determines ATG12 protein degradation via the UPS

G9a was previously known to inhibit autophagy by generating repressive histone H3K9 methylation in the promoters of autophagy genes ^26, 28^. However, our studies indicated that the *G9a* mRNA level is not altered under stress conditions (EBSS and UNC0638) employed in this study. In addition, we did not observe any alterations in the *ATG12* mRNA level, suggesting that ATG12 is not regulated by G9a at the transcriptional stage. Therefore, the reduction of G9a and ATG12 levels does not result from transcriptional regulation upon autophagy induction.

Our in vitro methylation studies revealed that K72 ATG12 is specifically methylated by G9a (Fig. 3c; Supplementary Fig. 8e,f). Indeed, we were able to detect K72me1 in ATG12 inside cells using specific antibodies raised against a peptide containing K72me1 (Fig. 3d; Supplementary Fig. 9). Moreover, we have shown that K72me1 ATG12 leads to the degradation of ATG12 via the UPS pathway (Fig. 3f–h), and that K72R ATG12, a mutant resistant to G9a-mediated methylation, reconstituted in ATG12-defincient MEF cells promotes autophagy (Fig. 7b). Therefore, methylation of K72 in ATG12 by G9a determines onset of autophagy. When K71 in ATG12 was replaced with R71 thereby generating K71R ATG12, a consensus G9a methylation site, R_71_K_72_ was generated. Thus, K71R ATG12 was hyper-methylated relative to WT ATG12 (Supplementary Fig. 8). Our findings were corroborated by the observation that WT ATG12 and K71R ATG12 underwent methylation at K72, whereas K72R ATG12 was not methylated (Fig. 3). Surprisingly, our Western blot data revealed that K71R ATG12 was less mono-methylated at K72 than WT ATG12. As reported for di-methylated LIG1 that is gradually depleted as tri-methylation of LIG1 proceeds via GLP ^19^. K72me1 in K71R ATG12 readily transitions to K72me2 by G9a, leading to the depletion of K72me1 Thus, we speculate that G9a/GLP prefer K72me1 ATG12 over WT ATG12 for methylation.

K90 in ATG12 is not a cognate G9a methylation site (Supplementary Fig. 8d); however, in the presence of the K71R (R_71_K_72_) mutation, G9a seems to recognize an additional methylation site, R_89_K_90_, as a cognate G9a methylation site (Supplementary Fig. 8f). This effect is reminiscent of β-catenin phosphorylation by a dual-kinase mechanism; phosphorylation at S35 by the CK1 kinase primes β-catenin for phosphorylation by GSK3β at S33, S37, and T41 ^67^. In a similar manner, K72 methylation might have primed R_89_K_90_ for additional methylation in the K71R/K89R ATG12 mutant. Furthermore, the K71R/K89R ATG12 mutant, which has two consensus G9a methylation sites (R_71_K_72_ and R_89_K_90_), is highly methylated (Supplementary Fig. 8f) and more ubiquitinated (Supplementary Fig. 11b) than other ATG12 mutants. Collectively, K72 residue in ATG12 is a methyl degron for ATG12 protein degradation by the UPS. Thus, the methylation status of K72 residue in ATG12 determines autophagy initiation.

### ATG12 methylation status is critical for autophagy induction and cell survival

Autophagy is intimately connected to apoptosis, and does not only inhibit apoptosis but also can promote cell death program depending on cell types and stress pathways ^68, 69^. Moreover, the same regulators such as ATG12 and Bcl2 are involved in both the processes ^70^. In this study, consistent with accumulating evidence that autophagy mostly inhibits apoptosis, we showed that K72R ATG12, a methylation resistant mutant, exhibits a significantly reduced apoptotic response compared with WT ATG12 and K71R ATG12, a hyper methylated mutant (Fig. 7c). However, recent studies have suggested a more complex relationship between autophagy and apoptosis in that ATG12 promotes mitochondrial apoptosis by forming the ATG12-ATG3 conjugate ^71^ and by interacting with anti-apoptosis Bcl2 ^70^. The reason for the discrepancy is not clear and warrants further studies, but it could be due to cell lines and apoptotic stimuli. In addition, activation of pro-autophagy initiation program by a hyper stable K72R ATG12 could have significantly raised a threshold for apoptosis. Taken together, G9a/GLP regulation of the ATG12 protein could be a promising therapeutic target for certain types of cancer resistant to apoptosis.

In summary, we here propose a novel mechanism (Fig. 7d) whereby G9a/GLP inhibit autophagy by directly methylating ATG12, which subsequently undergoes ubiquitination and degradation under non-stress conditions. However, the Ca^2+^-activated calpain system downregulates G9a/GLP proteins under stress conditions. This promotes an accumulation of the ATG12 protein and the formation of the ATG12-ATG5 conjugate and LC3II and subsequently expedites autophagosome formation.

## Methods

### Cell culture

ATG12+/+ and ATG12−/− MEFs were generously given to us by Dr. Jayanta Debnath (University of California San Francisco, CA, USA) ^72^. Wild type and ATG5-null MEFs were kindly provided by Dr. Noboru Mizushima (Tokyo Medical and Dental University, Tokyo, Japan) ^73^. MCF7, Huh7, Hela, U2OS, 293T, and MEF cells were cultured in DMEM supplemented with 10% fetal bovine serum (FBS) and 1× penicillin/streptomycin. Mouse embryonic stem cell lines (wild type TT2 and 248-5 cells that do not express both G9a and GLP) were obtained from Dr. Makoto Tachibana (Kyoto University, Japan) and cultured in DMEM containing 15% FBS, 10^3^ unit/mL LIF (Millipore), 0.1 mM β-mercaptoethanol (Gibco), 1% non-essential amino acids (Gibco), 1× penicillin/streptomycin, and 1× GlutaMax (Gibco).

### DNA construction and DNA and siRNA transfection

Amplicons encoding human calpain 1, calpain 2, CAST, ATG12, ATG5, ATG3, LC3B, JMJD2C, and G9a were digested with SalI and NotI and cloned into a SalI/NotI-digested N-terminal GST, C-terminal GST, N-terminal 5×Myc, C-terminal 3×HA, or C-terminal GFP vector. Lysines 71, 72, 89, and 90 of human ATG12 were mutagenized into arginine using site-directed mutagenesis following the manufacturer’s instructions (Agilent Technologies, QuikChange II XL site-directed mutagenesis kit). The plasmids encoding the indicated proteins were transfected into 293T cells using polyethylenimine (PEI), as described earlier ^60^. Lipofectamine 2000 was used for all transfection experiments with the exception of 293T cells. siRNAs were obtained from Ambion as follows: EHMT1/GLP (siRNA ID s534601); EHMT2/G9a (siRNA ID s21469); KDM4C/JMJD2C (siRNA ID s22990); and negative control No. 2 siRNA (# 4390846).

### Antibodies and reagents

The antibodies used in this study were as follows: anti-FLAG (Sigma, F1084); anti-myc, anti-Ub, anti-GFP, anti-HA, anti-GAPDH, and anti-JMJD2C (Santa Cruz, sc-40, sc-8017, sc-9996, sc-7392, sc-47724, and sc-104949, respectively); anti-GST (GE Healthcare, 27-4577-01); anti-G9a (Millipore, 09-071); anti-GLP (R&D, PP-B0422-00); anti-β-Actin (Thermo Fisher, MA1-140), anti-LC3B (Novus, NB100-2220); anti-ATG5, anti-ATG12, anti-GAPDH, anti-DCAF1/VPRBP, p62, and p70 S6 kinase (Cell Signaling, #12994, #4180, #2118, #14966, #5114, and #2708, respectively); anti-H3K9me2 and anti-histone H3 (Abcam, ab1220 and ab1791); anti-ATG12 K72me1 (Creative Biolab Inc.); anti-rabbit IgG, light chain specific HRP conjugate (Jackson ImmunoResearch, 211-032-171); and anti-mouse IgG linked HRP and anti-rabbit IgG linked HRP (Cell Signaling, #7076 and #7074). Inhibitors and their final concentrations were as follows: UNC0638 (2-10 μM, Sigma, U4885), JIB 04 (1 μM, Axon, NSC 693627), ML 324 (1 μM, Axon, 2081), cycloheximide (50 μg/mL, Sigma, C4859), MG132 (10 μM, Santa Cruz, sc-351846), A23187 (1 μM, Santa Cruz, sc-3591), chloroquine (50 μM, Sigma, C6628), Torin1 (1 μM, Santa Cruz, sc-396760), BAPTA/AM (10 μM, Santa Cruz, sc-202488), Calpeptin (2-10 μM, Santa Cruz, sc-202516), and bafilomycin A1 (20 nM, Santa Cruz, sc-201550). Autophagy was induced by starvation in Earle’s Balanced Salt Solution (EBSS, Sigma, E6132) (116 mM NaCl, 5 mM KCl, 1.8 mM CaCl_2_·2H_2_O, 0.4 mM MgSO_4_, 1 mM NaH_2_PO_4_, 5 mM Glucose, 3 µM Phenol red, pH 7.6), in HEPES-based starvation media (140 mM NaCl, 1 mM CaCl_2_, 1 mM MgCl_2_, 5 mM Glucose, 20 mM HEPES pH 7.4, and 1% BSA) ^36^ and by inhibiting mTOR with 1 μM Torin1 for the indicated time.

### Microscopy

Confocal images were acquired using a Zeiss LSM700 confocal microscope equipped with a 40× or 60× water objective and digitally captured using LSM software. Fluorescence images were obtained using a Leica DM5000 B upright microscope. To measure the mean fluorescent intensity (MFI), the area of interest was outlined at the indicated magnifications based on the fluorescent signal and using a Leica upright fluorescence microscope. After setting a manual threshold and background subtraction, fluorescent intensity was quantified using ImageJ software.

### Immunoblots and co-immunoprecipitation

For Western blot analyses, expression plasmids were transfected into 293T, Hela, U2OS, or ATG12−/− MEF cells using PEI or Lipofectamine 2000 according to the manufacturer’s instructions (Invitrogen). Two days after transfection, the cells were washed with PBS and lysed with RIPA buffer (20 mM Tris-HCl pH 7.5, 150 mM NaCl, 1 mM Na_2_EDTA, 1 mM EGTA, 1% NP-40). The cell lysates were collected and used for Western blotting and other experiments. Protein samples (20 µg) from each lysate were subjected to SDS-polyacrylamide gel electrophoresis (PAGE), transferred to a PVDF membrane, and probed with specific primary antibodies. Proteins of interest were detected using an HRP-conjugated secondary antibody and ECL substrates (Amersham) and visualized with the ChemiDoc MP system (Bio-Rad). Rabbit IgG heavy chains of anti-ATG12 antibody overlap with the ATG12-ATG5 conjugate. To overcome this limitation, an anti-rabbit IgG secondary antibody-conjugated HRP specific for light chains (Jackson ImmunoResearch, 211-032-171) was used after incubating the membrane with anti-ATG12 K72me1 antibody. The intensity of each protein band was normalized using GAPDH or β-Actin for relative quantitation in the ImageJ program. To identify endogenous protein–protein interactions, cell lysates were sonicated and treated with DNase I to remove genomic DNA contamination and then incubated with an anti-ATG5 or anti-ATG12 antibody (5 μg of either). Rabbit IgG (5 µg) was added to the lysates as a control. Unless otherwise indicated, all the experiments were repeated at least three times.

### Subcellular fractionation assay

The cytoplasmic and nuclear fractions were isolated with an EpiQuik nuclear extraction kit (EpiGentek, OP-0002) according to the manufacturer’s protocol. Briefly, cells were washed with PBS and collected by centrifugation. The cell pellet was resuspended in 100 µl of NE1 solution, incubated on ice for 10 min, and then centrifuged at 12,000 rpm for 1 min at 4°C. The supernatant was retained as the cytoplasmic fraction. The pellet was incubated in 2 volumes of NE2 containing DTT and PIC on ice for 15 min with vortexing every 3 min. The extract was further sonicated three times for 10 seconds each time to increase the nuclear protein extraction. After centrifuging the suspension for 10 min at 14,000 rpm and 4°C, the supernatant was retained as the nuclear fraction.

### GST pull-down and immunoprecipitation assay

For the GST pull-down assays, we used a mammalian GST-fusion protein expression system. GST-fusion constructs were transfected into 293T cells using PEI reagent. Two days after transfection, the cells were lysed with Pierce IP lysis buffer (25 mM Tris-HCl pH 7.4, 150 mM NaCl, 1% NP-40, 1 mM EDTA, and 5% glycerol) containing a protease inhibitor cocktail (Roche,11 697 498 001). The cell lysates were further sonicated on ice for 10 seconds, and the supernatant was collected by centrifugation at 13,000 rpm at 4°C for 20 min. Glutathione Sepharose 4B was added to the clarified lysates and incubated at 4°C overnight. For the immunoprecipitation assay, protein A/G PLUS-Agarose beads (Santa Cruz, sc-2003) were added to the clarified lysates and incubated at 4°C for 2 hours. The beads were removed by centrifugation for 5 min at 13,000 rpm. Then, specific primary antibody (5 μg) was added to the bead-free lysates and incubated at 4°C for 2 hours. After this step, the beads were added again and incubated at 4°C overnight for immunoprecipitation. The next day, the beads were collected by centrifugation and washed three times with RIPA buffer for 15 min each time. Thereafter, 1× SDS sample buffer was added to the pellets, and the samples were analyzed by SDS-PAGE and Western blots. To detect ubiquitination, GST-fusion constructs and HA-Ub were transfected into 293T or Hela cells. The remainder of the GST pull-down assays was conducted as described above. Ubiquitination was detected using an anti-HA (Santa Cruz, sc-7392) or anti-Ub (Santa Cruz, sc-8017) antibody.

### *In vitro* calpain 1 cleavage assay

Recombinant Flag-tagged human G9a (Active Motif, 31410), human calpain 1 (Novus, P4041), and calpain S1 (Abcam, ab180298) were incubated at 37°C for 10 min in calpain buffer (20 mM HEPES pH 7.5, 50 mM KCl, 2 mM MgCl_2_, 5 mM CaCl_2_, 1 mM DTT). The cleavage reaction was terminated by the addition of an equal volume of 2× SDS sample buffer and boiling at 95°C for 10 min.

### *In vitro* G9a methyltransferase assays using GST-fusion proteins

293T cells were transfected with plasmid to express GST alone or GST-ATG12, GST-ATG5, GST-LC3B, and GST-G9a (SET). Two days after transfection, the transfected cells were harvested and lysed in Pierce IP lysis buffer. The cell lysates were centrifuged for 10 min at 13,000 rpm and 4°C. The supernatants were collected and incubated with glutathione Sepharose 4B beads for 16 hours at 4°C. The beads were collected by centrifugation and washed three times with RIPA buffer. To purify the GST fusion proteins, Sepharose beads bound to GST fusion proteins were incubated with reduced glutathione (10 mM) for 30 min. The beads were centrifuged at 5,000 rpm for 10 min at 4°C, and the supernatants containing GST fusion proteins and glutathione were carefully collected and subjected to dialysis. Finally, the GST fusion proteins were mixed together and incubated in methylation buffer (50 mM Tris-HCl pH 8.8, 5 mM MgCl_2_, and 4 mM DTT) containing S-adenosyl-l-[methyl-^3^H] methionine (85 Ci/mmol from a 0.5 mCi/mL stock solution, Perkin-Elmer) for 6 hours at room temperature. The reaction was stopped by the addition of 2× SDS sample buffer and subjected to SDS-PAGE followed by transfer onto a PVDF membrane. The ^3^H radiographic signal was acquired with a storage phosphor screen (GE, BAS-IP TR2025 E).

### *In*-gel digestion

Upon separation by SDS-PAGE and staining with Coomassie Blue, the protein bands corresponding to the ATG12 of the PAGE gel excised, destained in 50 mM ammonium bicarbonate with 50% acetonitrile (ACN), dehydrated with 100% ACN and then dried in a vacuum evaporator. Gel pieces were rehydrated with 10 mM dithiothreitol and incubated 1 hour at 56 °C followed by 55 mM iodoacetamide and incubated 1 hour at 25 °C. After thiol blocking, Glu-C (Sigma-Aldrich, St. Louis, MO, USA) digest was performed. Initially, the gel pieces rehydrated with 12.5 ng/μL Glu-C in 25 mM ammonium bicarbonate and incubated at 4 °C for 2 hours then discarded supernatants and replenished with 50 mM ammonium bicarbonate. Digestion was performed at 37 °C for 16 hours. Peptides were extracted with 25 mM ammonium bicarbonate in 50% ACN, subsequently 0.5 % trifluoroacetic acid (TFA) in 50% ACN and dried in a vacuum evaporator

### Liquid chromatography and tandem mass spectrometry

Dried peptide samples were reconstituted in 7 µL of 0.1% formic acid, and an aliquot containing 5 μL was injected from a cooled (10 °C) autosampler into a reversed-phase Peprosil-Pur 120 C18 AQ, 5 µm (Dr. Maisch GmbH, Ammberbuch-Entringen, Germany) column (15 cm × 75 μm, packed in-house) on an Eksigent nanoLC 2D system at a flow rate of 300 nL/min. Prior to use, the column was equilibrated with 95% buffer A (0.1% formic acid in water) and 5% buffer B (0.1% formic acid in acetonitrile). The peptides were eluted with a linear gradient from 5% to 30% buffer B over 70 min and 30% to 70% buffer B over 5 min followed by an organic wash and aqueous re-equilibration at a flow rate of 300 nL/min with a total run time of 120 minutes. The HPLC system was coupled to an LTQ-Orbitrap XL mass spectrometer (ThermoFisher Scientific, Bremen, Germany) operated in a data-dependent acquisition (DDA) mode. Full scans (m/z 300–1800) were acquired at a resolution of 60,000 using an AGC target value of 1e6 and a maximum ion injection time of 10 ms. Tandem mass spectra were generated for up to 5 precursors by collision induced dissociation (CID) in ion-trap using a normalized collision energy of 35%. The dynamic exclusion was set to 60 s. Fragment ions were detected at a normal scan mode using an AGC target value of 1e5 and a maximum ion injection time of 500 ms. Source ionization parameters were as follow: spray voltage, 1.9 kV; capillary temperature, 275 °C.

### Quantitative real time polymerase chain reaction (qRT-PCR)

Total RNA was isolated from Hela cells (n = 3) subjected to EBSS-induced autophagy and treatment with 5 μM UNC0638 for 12 hours using TRIzol reagent (Invitrogen) according to the manufacturer’s instructions. cDNA was synthesized from 1 μg of total RNA with random primers using an iScript cDNA synthesis kit (Bio-Rad) and analyzed with a Bio-Rad CFX96 real time system. All the experiments were performed at least three times in duplicate. The relative expression levels of the mRNAs were assessed and normalized to GAPDH expression.

*G9a*-5: TTCTGGGACATCAAAAGCAA, *G9a*-3: TGTGGTCCGTTCTCATGTGT

*JMJD2C*-5: TTCCACAGTCAGGGAAGGAC, *JMJD2C*-3: TCATTTCACAAAGGCCAACA

*ATG10*-5: CTTGATCCTGGGAGTTCGAG, *ATG10*-3: TGGCACAGTCTCAGCTCACT

*ATG5*-5: GGCAGCCAGTCTTTTGAAGT, *ATG5*-3: GACAGCCAGCTAGCTCACAA

*ATG7*-5: CAGCGTAACGATGGTCTCAA, *ATG7*-3: AATTGGCACTGGGTGAAAAG

*ATG12*-5: CCATTCTTTGAAGATGCCTTG, *ATG12*-3: CATGGCTCTGAGGCAGTATG

*LC3B*-5: CAGCGGGAGAAACACAAAAT, *LC3B*-3: ATGCACTTGGTGTGGGAGA

*WIPI1*-5: AATCTTGTGCCGTGGAAATC, *WIPI1*-3: GGGAAGCTTGGTGAGGAGTT

*GAPDH*-5: TTGAGGTCAATGAAGGGGTC, *GAPDH*-3: GAGGTGAAGGTCGGAGTCA

### Statistical analysis

Data are presented as the mean ± SEM of at least three independent experiments. Student’s t-tests were used for parametric data analyses to compare groups using GraphPad Prism (version 6.0) software (GraphPad Software, Inc., San Diego, CA). In the figures, ∗p < 0.05, ∗∗p < 0.01, and ∗∗∗p < 0.001 denote statistical significance.

## Acknowledgments

We are grateful to Dr. Jayanta Debnath (The University of California, San Francisco, CA, USA) for the ATG12+/+ and ATG12−/− MEFs, Dr. Noboru Mizushima (Tokyo Medical and Dental University, Tokyo, Japan) for the wild-type and Atg5 knockout MEFs, and Dr. Makoto Tachibana (Kyoto University, Kyoto, Japan) for the mouse TT2 and 248-5 embryonic stem cells. We also appreciate Dr. Wen-Xing Ding (The University of Kansas Medical Center, Kansas City, KS, USA) for his valuable comments and suggestions. This research was supported by grants 2017R1A5A2015395 (The Medical Research Center to S.L.), 2017M3A9B4062401 (to S.L.), NRF-2017R1A2B2002220 (to S.L.), and NRF-2015R1D1A1A01060986 (to C.K.) from the National Research Foundation of Korea (NRF) of the Ministry of Science and ICT and the Ministry of Education, Republic of Korea. The Intramural Research Program (Theragnosis) of KIST also supported this study.

## Author Contributions

C.K. conceived the idea and planned the project; C.K., K.P., S.J. and C.L. performed experiments; C. K., K.P., S.J., C.L. and S.L. analyzed the data; C.K., K.P., and S.L. wrote the manuscript; all authors discussed the results and approved this manuscript.

## SUPPLEMENTARY INFORMATION

### SUPPLEMENTARY FIGURE LEGENDS

**Supplementary Figure 1.**
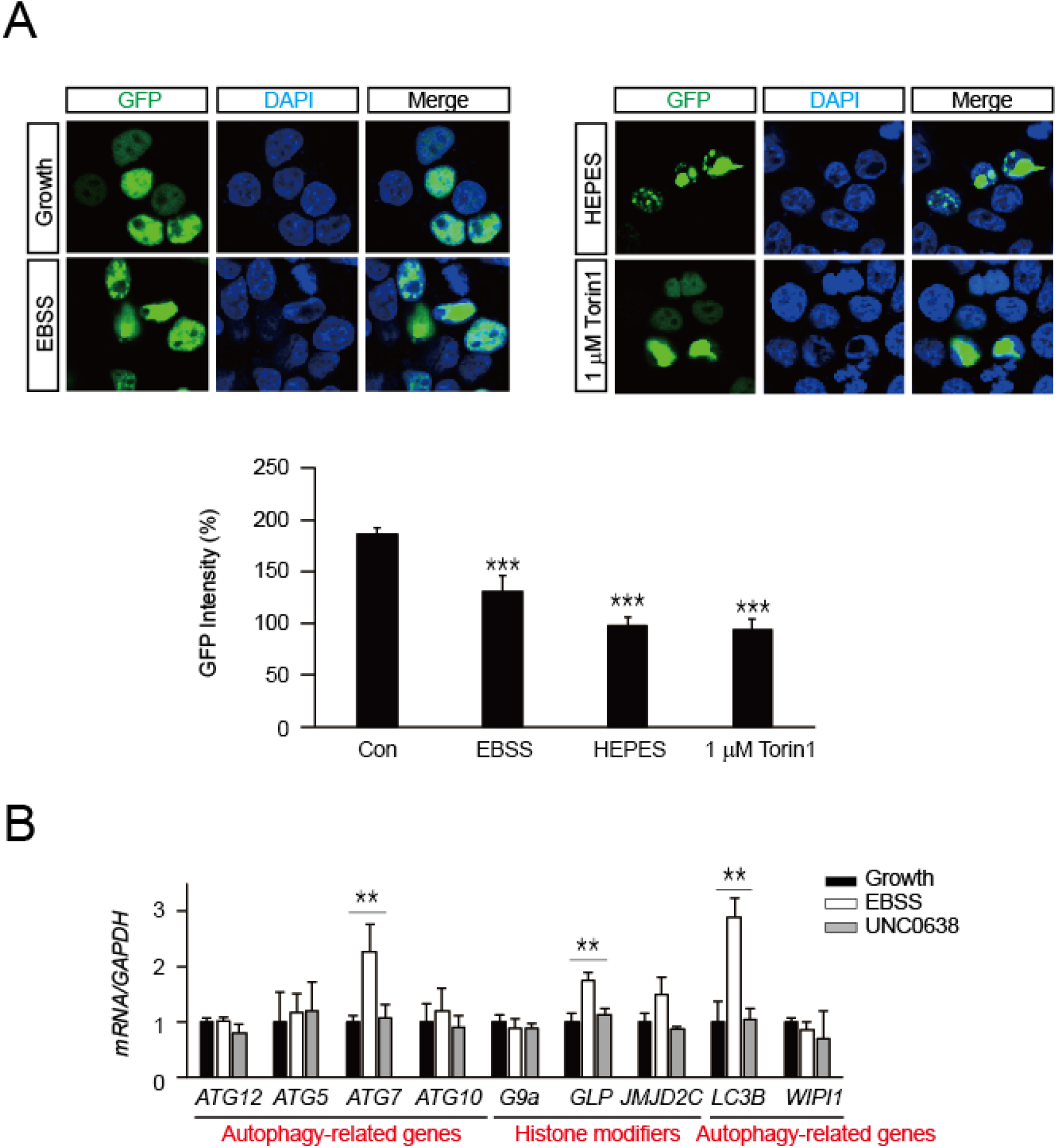
G9a protein downregulation during autophagy involves post-translational regulation. (a) G9a-GFP is downregulated during autophagy induced by serum starvation (EBSS and HEPES) and mTOR inhibition (1 µM Torin1) in 293T cells. GFP fluorescence intensity was quantified as described in Figure 1B (n = 3 [50–90 per each group]; ***p < 0.001; error bars, ±SEM). (b) Autophagy induced by starvation in EBSS medium and G9a inhibition (5 µM UNC0638) for 12 hours did not alter the *G9a* mRNA level but increased the *GLP* and *JMJD2C* mRNA levels in Hela cells. Experiments were repeated three times, and student’s t tests were used for a relative comparison between *GAPDH* and *mRNAs* levels (mean ± SEM of n = 3 replicates; **p < 0.01).

**Supplementary Figure 2.**
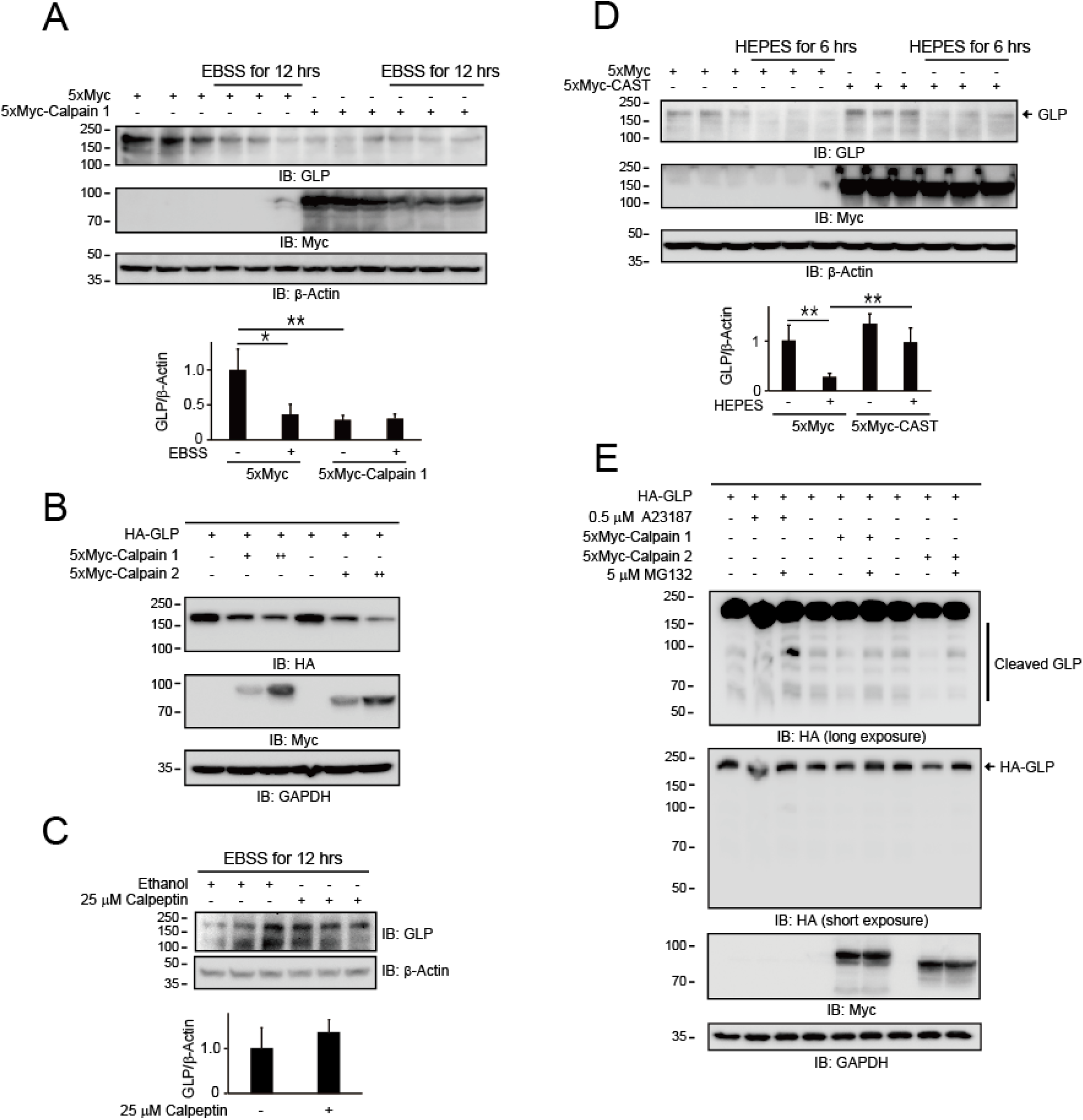
GLP, a member of the G9a family, is cleaved by calpains. (a) Calpain 1 expression downregulated the endogenous GLP protein levels in 293T cells. The samples were manipulated as described in Figure 1D (mean ± SEM of n = 3 replicates; *p < 0.05; **p < 0.01). (b) HA-GLP was cleaved and degraded by calpain 1 and 2 expression in 293T cells. (c) A calpain inhibitor, calpeptin, elevated the endogenous GLP protein level in 293T cells under starvation in EBSS medium (mean ± SEM of n = 3 replicates). NS no significant difference (Student’s *t*-test). (d) Expression of CAST/Calpastatin, a calpain inhibitory protein, in Hela cells significantly stabilized endogenous GLP under starvation in HEPES medium (mean ± SEM of n = 3 replicates; **p < 0.01). (e) Cleaved HA-GLP was degraded by the UPS pathway. MG132, a proteasome inhibitor, led to the accumulation of the cleaved GLP intermediates produced by elevated Ca^2+^ influx (0.5 µM A23187) and the expression of calpains 1 and 2 in 293T cells.

**Supplementary Figure 3.**
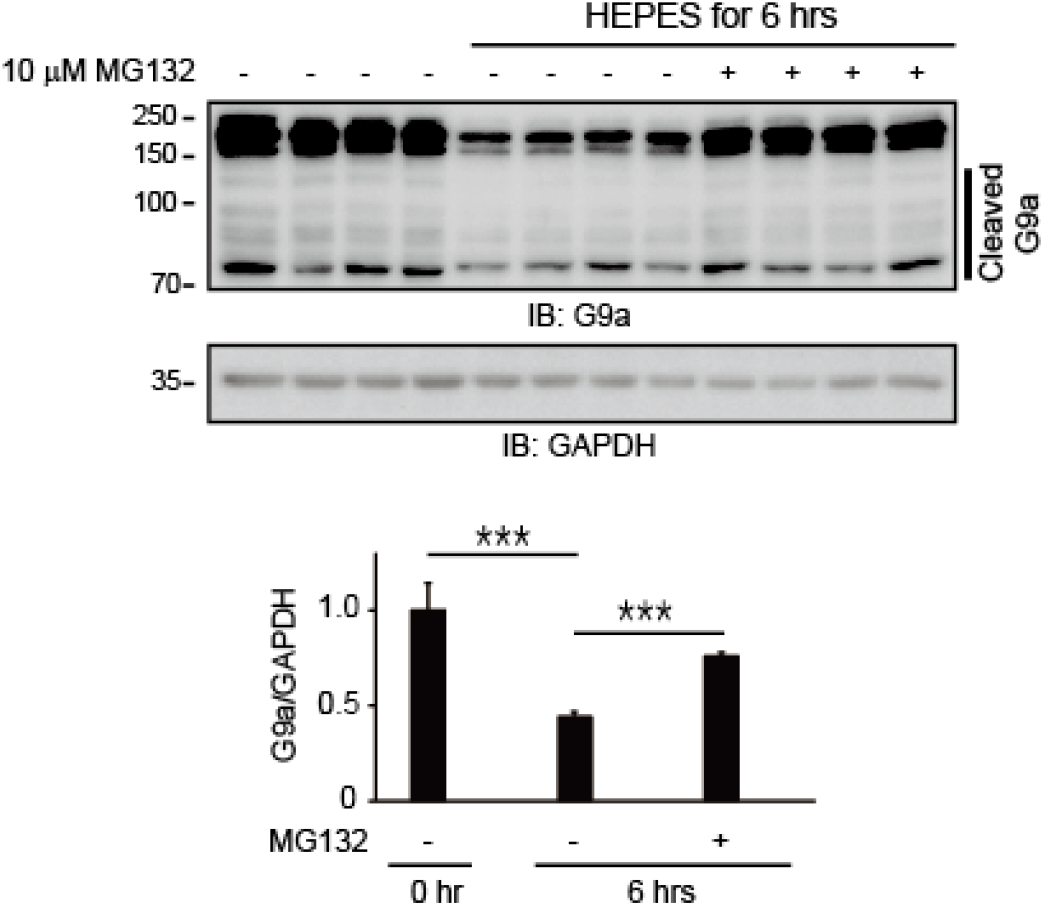
Endogenous G9a undergoes degradation via the UPS during autophagy. Autophagy induction by serum starvation (HEPES) diminished the endogenous G9a protein level in Hela cells. In the presence of MG132, a proteasome inhibitor, endogenous G9a degradation was significantly reduced under starvation in HEPES medium, indicating that G9a degradation was facilitated by the UPS (mean ± SEM of n = 4 replicates; ***p < 0.001).

**Supplementary Figure 4.**
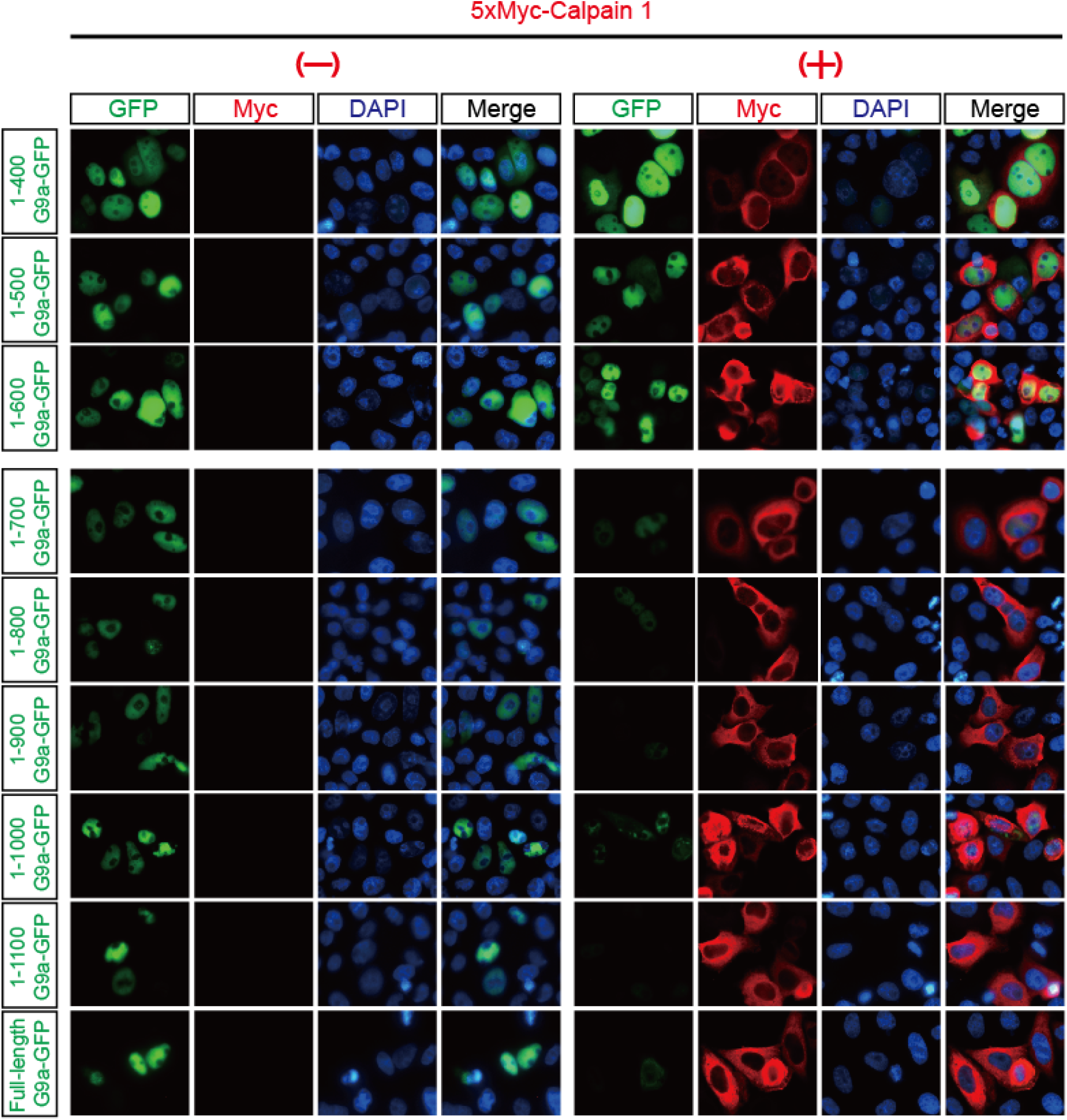
A calpain cleavage site in G9a is located between aa 600 and aa 700. Fading fluorescence intensity from the truncation G9a mutants fused to a C-terminal GFP indicates that calpain 1 cleaved between aa 600 and aa 700 in G9a. Constructs expressing truncated G9a-GFP with or without 5xMyc-calpain 1 were transfected into Hela cells, and GFP signals were acquired using a Leica DM5000 B.

**Supplementary Figure 5.**
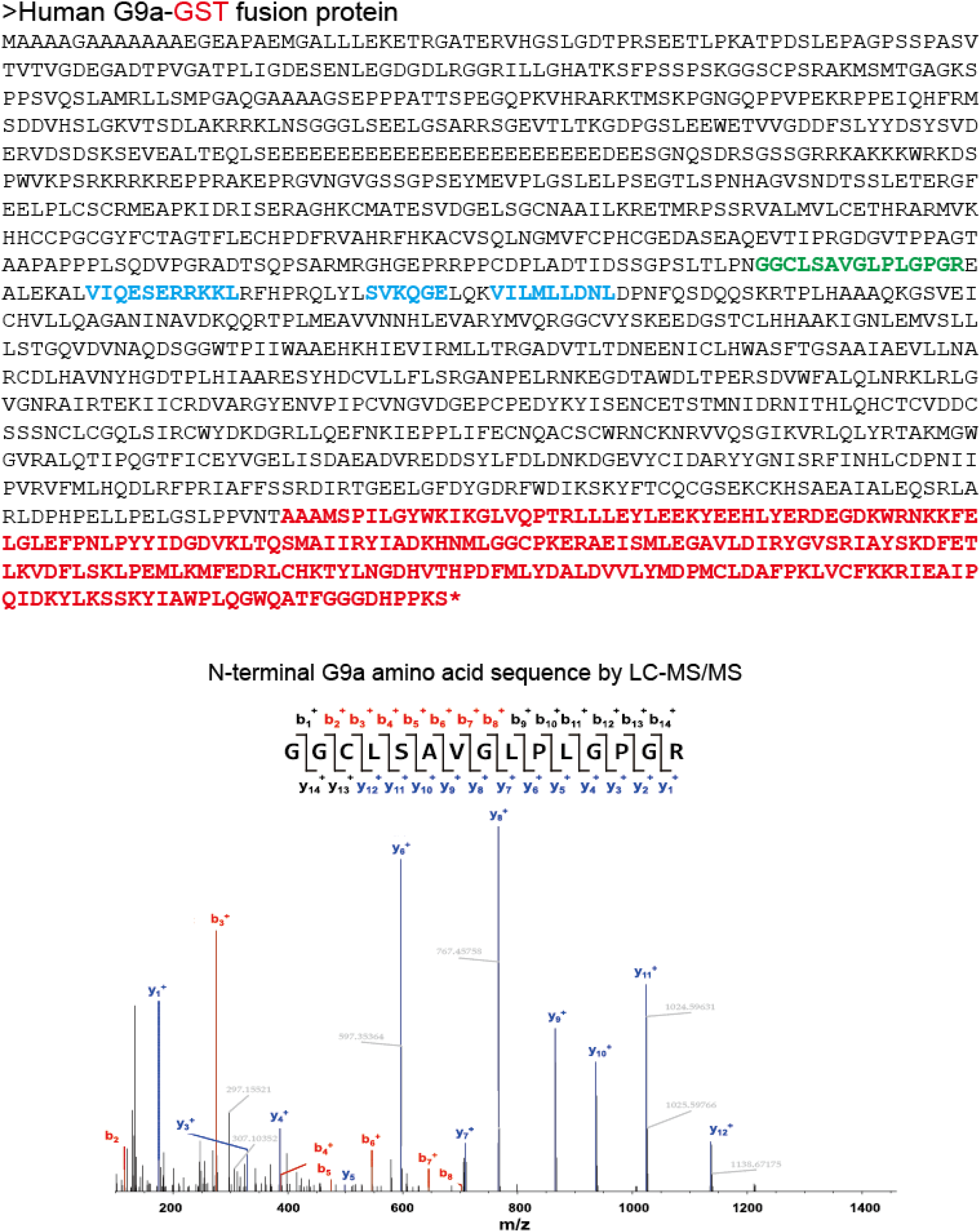
LC-MS/MS analysis confirms that the calpain cleavage site in G9a is located between aa 600 and aa 700. Peptide mapping by LC-MS/MS indicates that calpain 1 cleaved at the 615 position in G9a starting with a GGCLSAV sequence (green). In addition, LC-MS/MS data reveal three other calpain cleavage sites (blue). G9a-GST and 5xMyc-calpain 1 were transfected into 293T cells. The cleaved G9a-GST was purified using a glutathione Sepharose column as described in the STAR Methods and digested with trypsin prior to analysis with LC-MS/MS.

**Supplementary Figure 6.**
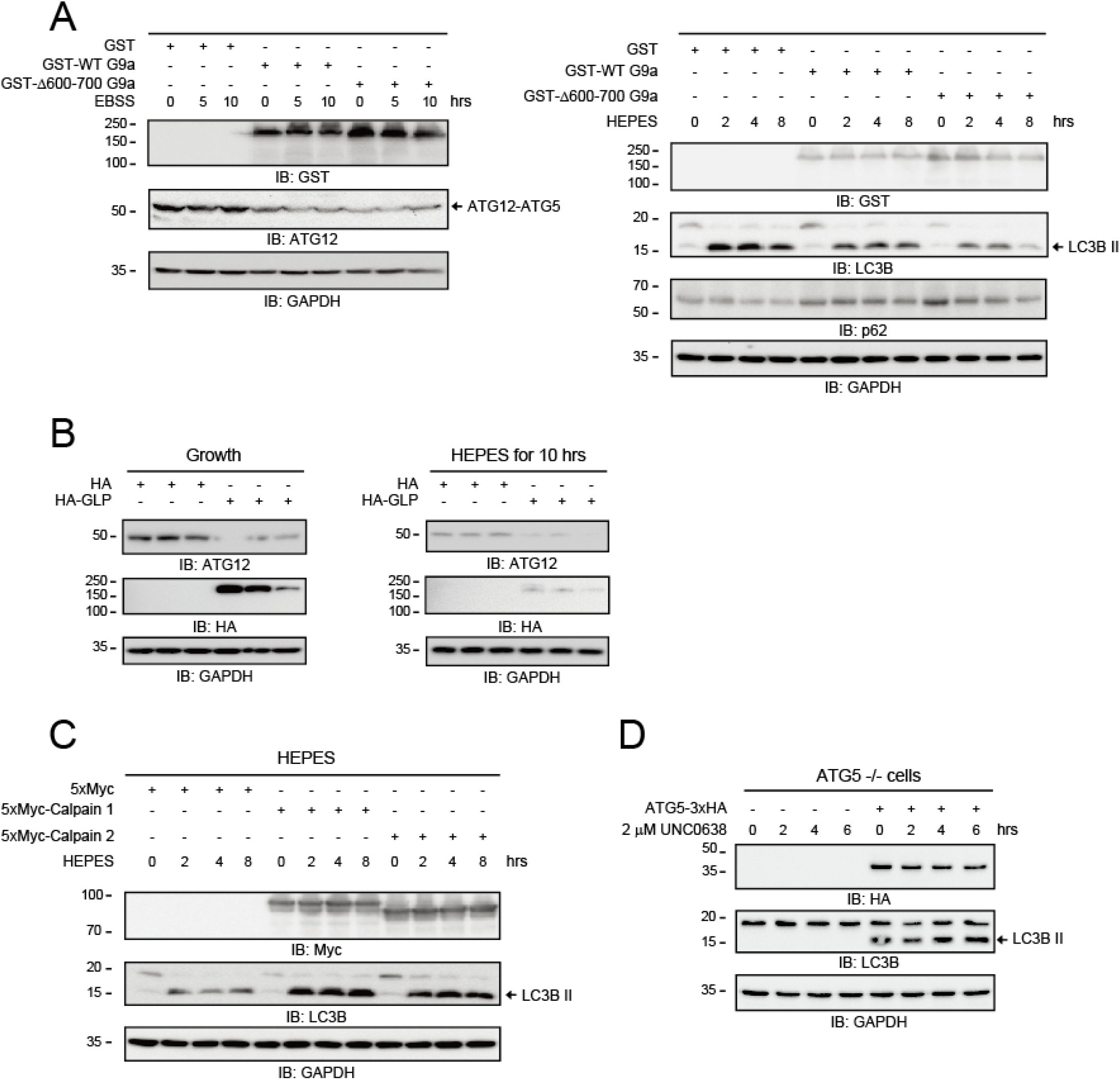
G9a inhibits autophagy induction by downregulating the ATG12-ATG5 conjugate level. (a) Both GST-WT G9a and GST-Δ600–700 G9a expression downregulated the ATG12-ATG5 conjugate level under starvation in EBSS medium. The expression of the calpain-resistant Δ600–700 G9a in 293T cells suppressed the formation of the ATG12-ATG5 conjugate and the conversion of LC3BI to LC3BII more noticeably than the expression of WT G9a while increasing the p62 level under starvation conditions. (b) HA-GLP expression in Hela cells also reduced the ATG12-ATG5 conjugate level under growth and starvation (HEPES) conditions. (c) The expression of calpains 1 and 2 promoted the conversion of LC3BI to LC3BII under starvation in HEPES medium. (d) Autophagy induction by G9a inhibition (2 µM UNC0638) required ATG5. Only when ATG5 was reconstituted in ATG5−/− MEFs, UNC0638-mediated LC3BII conversion occur. Thus, ATG12-ATG5 conjugate formation was necessary for UNC0638-induced autophagy.

**Supplementary Figure 7.**
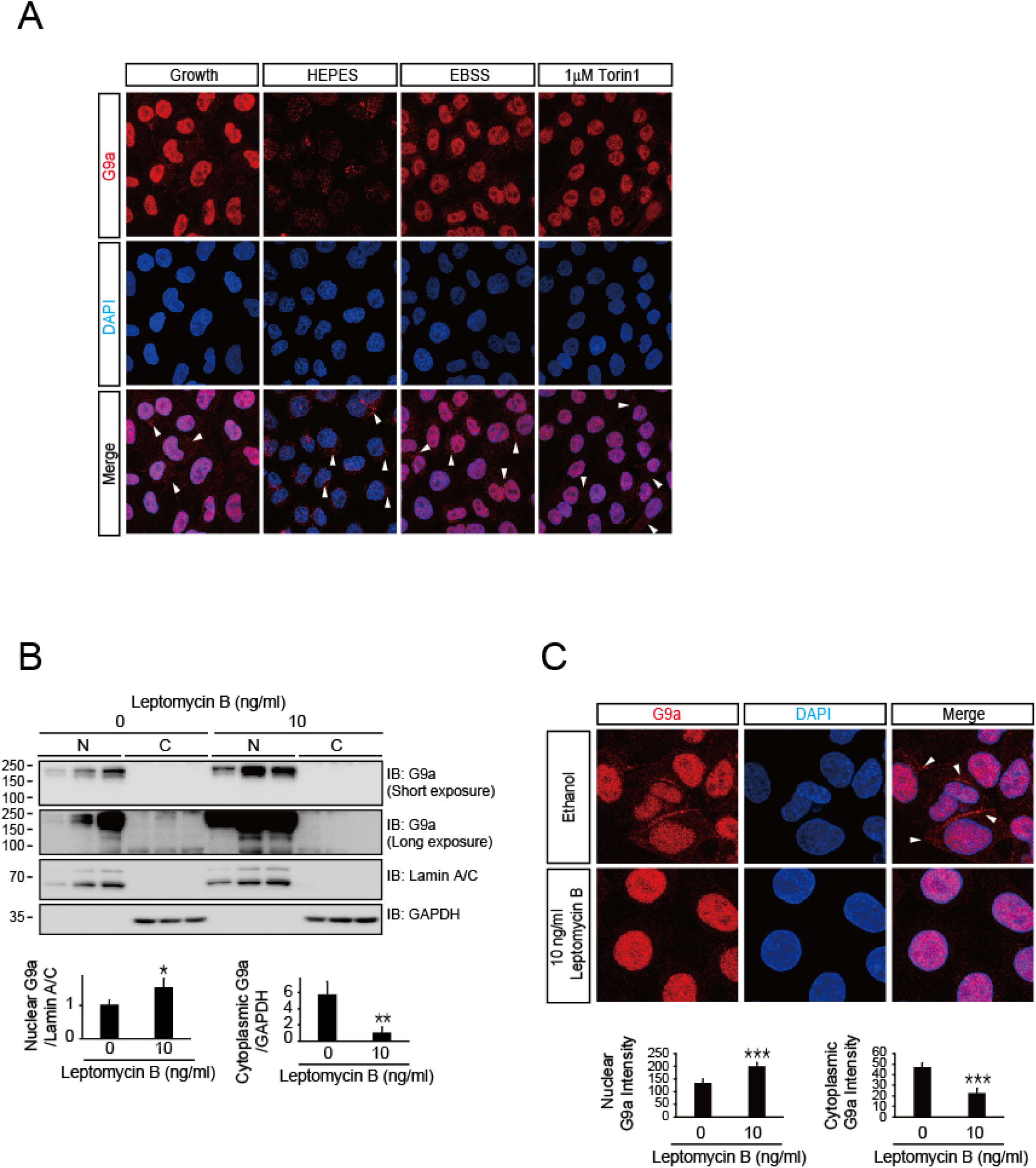
Endogenous G9a localizes in both cytoplasm and nucleus. (a) Confocal IF staining reveals that endogenous G9a was diminished and exhibited slightly enhanced localization in the cytoplasm under starvation (EBSS and HEPES) and mTOR inhibition (1 µM Torin1). The white arrow heads indicate cytoplasmic G9a proteins. (b) Treatment with Leptomycin B increased the nuclear G9a level and decreased the cytosolic G9a level. The samples were manipulated as described in Supplementary Figure7A. GAPDH and Lamin A/C were probed as cytoplasmic and nuclear fractionation controls, respectively. N, nuclear fractions; C, cytoplasmic fractions (mean ± SEM of n = 3 replicates; *p < 0.05; **p < 0.01). (c) Confocal IF imaging shows that Leptomycin B clearly decreased the cytosolic G9a level. The white arrow heads indicate cytoplasmic G9a proteins. The intensity of nuclear and cytosolic G9a protein was quantified using MFI of individual cells (n = 3 [30-50 per each group]; ***p < 0.001; error bars, ±SEM).

**Supplementary Figure 8.**
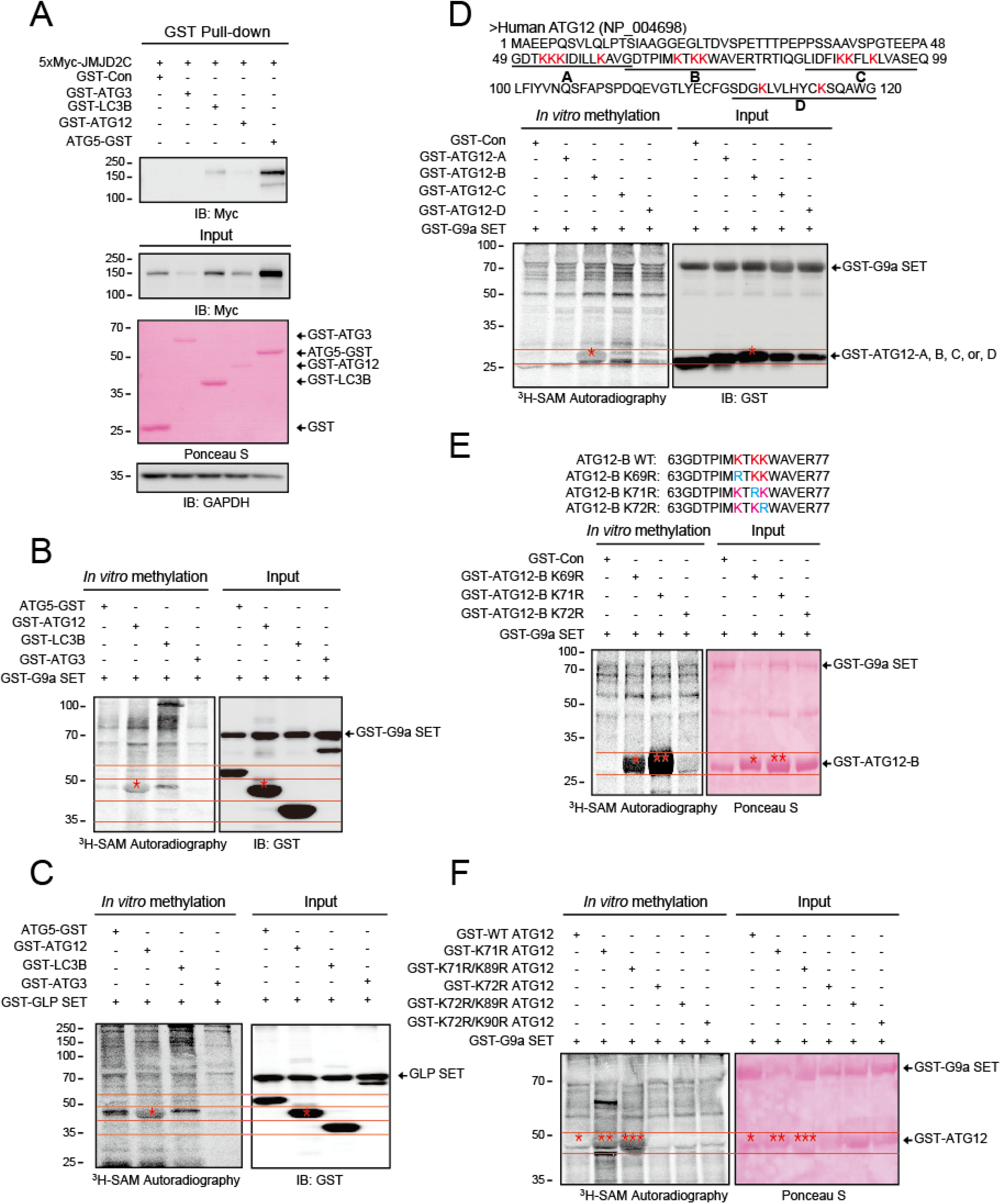
ATG12 methylated by GLP interacts with JMJD2C. (a) Pull-down assay shows that JMJD2C interacted with ATG12, ATG5, and LC3B in 293T cells. (b) A catalytic G9a SET methylated only ATG12 in *in vitro* methyltransferase assays. (c) The GST-GLP SET construct methylated ATG12 in an *in vitro* methyltransferase assay. (d) *In vitro* methyltransferase assays show that G9a methylated the K residues between aa 63 and aa 77 in the GST-ATG12-B protein. (e) K72 residue in ATG12 is a specific methylation site of G9a. Purified GST-Con and GST-ATG12-B fragment proteins containing K69R, K71R, or K72R were used for *in vitro* methyltransferase assays. (f) The methylation degree of full-length ATG12 mutants was enhanced by the presence of consensus G9a methylation sites. K71R ATG12 (R_71_K_72_) and K71R/K89R ATG12 (R_71_K_72_/R_89_K_90_) exhibited much higher levels of methylation than WT ATG12. Purified full-length GST-WT ATG12 and GST-ATG12 mutant proteins were used for *in vitro* methyltransferase assays. *, **, and *** mark peptide(s) methylated by G9a SET.

**Supplementary Figure 9.**
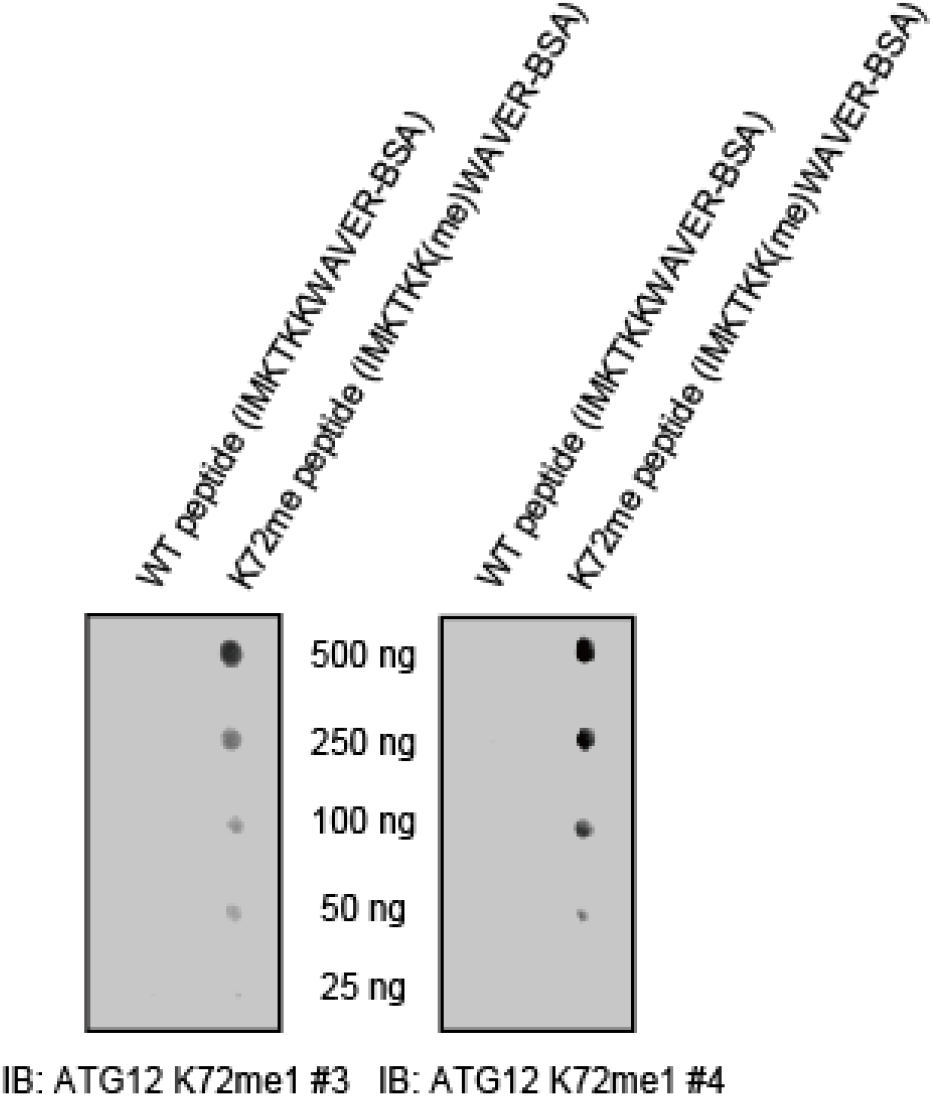
Validation of anti-ATG12 K72me1 antibodies. Dot blots confirm that polyclonal antibodies specifically recognized the K72me1 peptide. Antibody batch #4 was less sensitive than batch #3, but it was more specific.

**Supplementary Figure 10.**
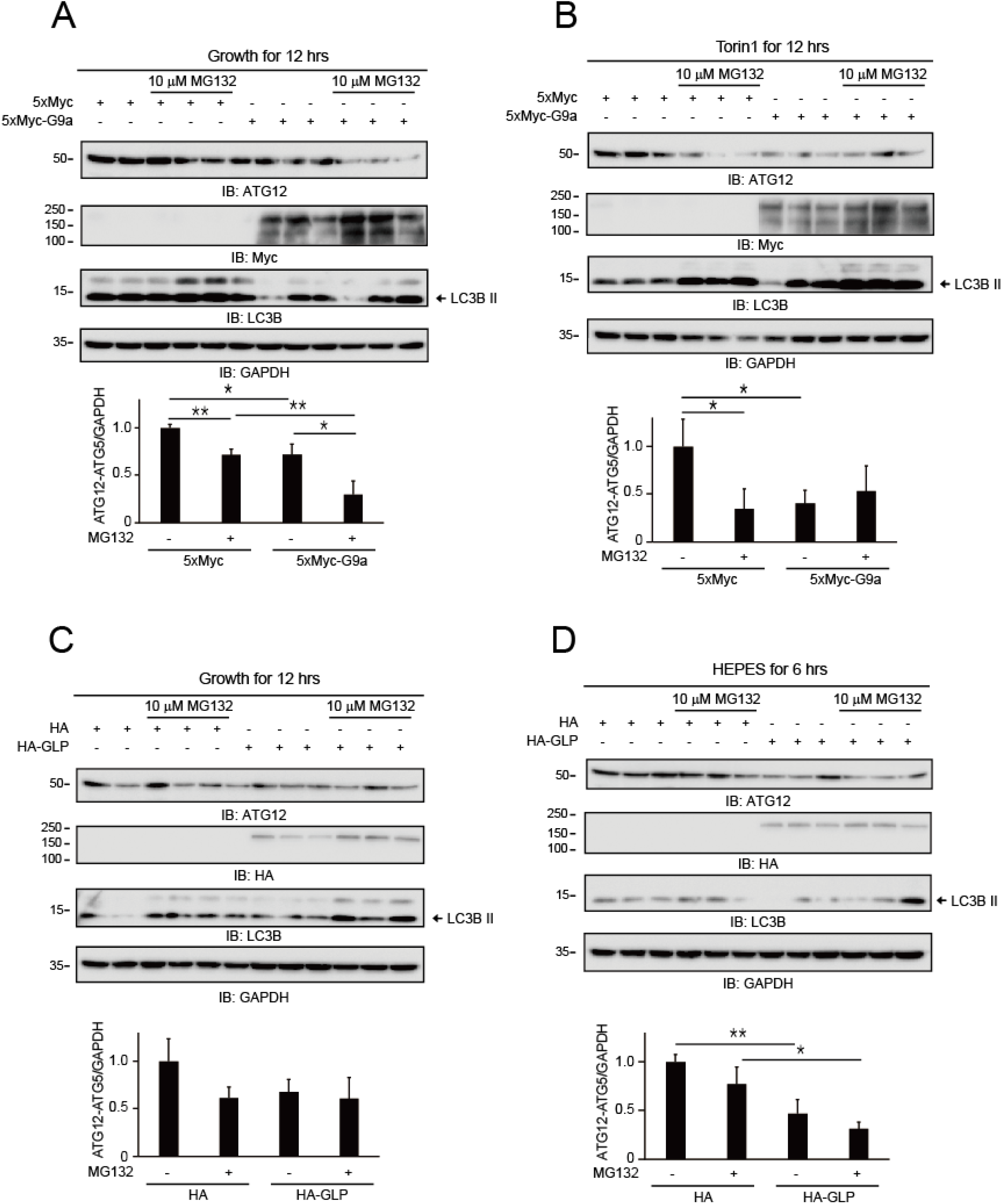
MG132 does not prevent G9a/GLP-induced reduction of the ATG12-ATG5 conjugate. G9a/GLP expression decreased the ATG12-ATG5 conjugate level in Hela cells, and MG132 did not prevent the reduction. On the contrary, MG132 reduced the ATG12-ATG5 conjugate level through a lysosome-mediated protein degradation pathway based on the increased LC3BII level under growth (**a** and **c**), mTOR inhibition (**b**), and HEPES starvation (**d**) conditions. Therefore, G9a/GLP expression did not accumulate the ATG12-ATG5 conjugate level in the presence of MG132 (mean ± SEM of n = 3 replicates; *p < 0.05; **p < 0.01).

**Supplementary Figure 11.**
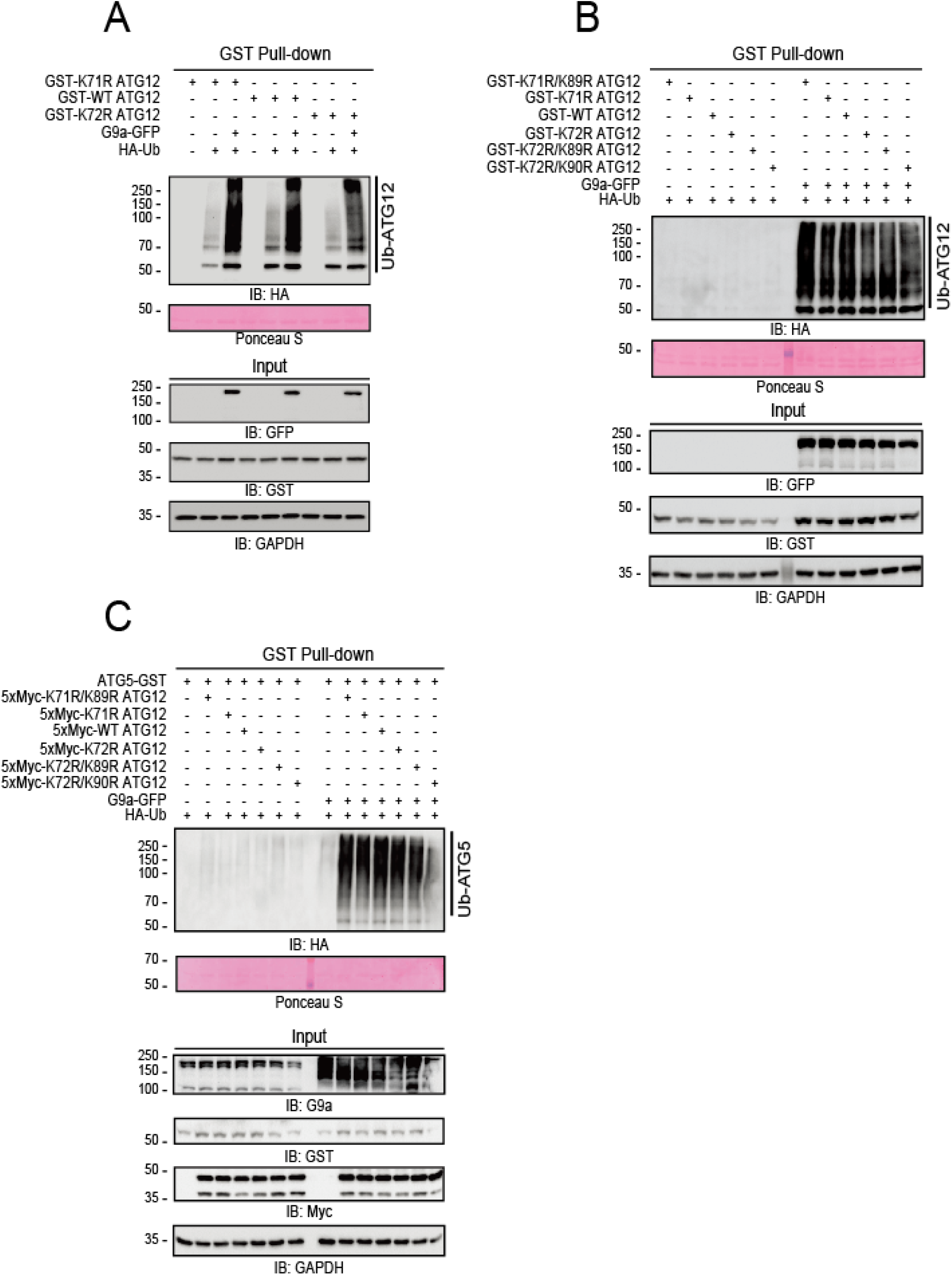
Methylation status of K72 residue in ATG12 is coupled to ATG12 protein degradation by the UPS. (a) G9a promoted the ubiquitination of ATG12 mutants in the following order: K71R ATG12 > WT ATG12 > K72R ATG12. GST pull-down assays were performed to determine the ubiquitination levels of the GST-ATG12 mutants. (b) Ubiquitination of the ATG12 mutants required G9a expression. The ubiquitination levels of the ATG12 mutants were ranked as follows: K71R/K89R ATG12 > K71R ATG12 > WT ATG12 > K72R ATG12 > K72R/K89R ATG12 > K72R/K90R ATG12. (c) ATG5 ubiquitination was determined by the degree of ATG12 methylation.

**Supplementary Figure 12.**
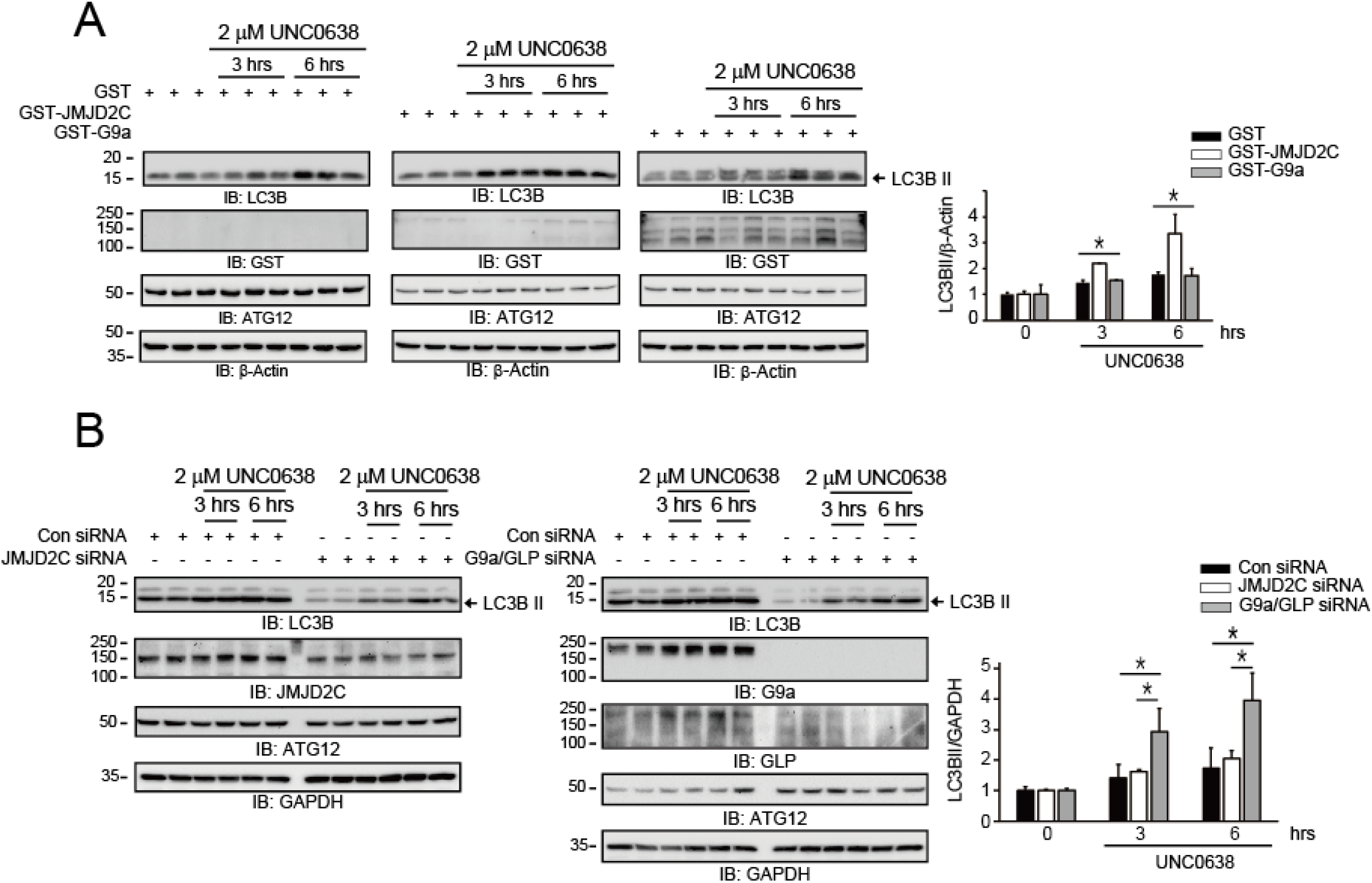
The endogenous ATG12-ATG5 conjugate level set by G9a/GLP and the JMJD2 family determines UNC0638-mediated LC3BII formation. (a) JMJD2C significantly augmented UNC0638-mediated autophagy induction in Hela cells. (b) Silencing both G9a and GLP expression significantly enhanced autophagy induction, whereas silencing JMJD2C expression had little effect compared with control siRNA in Hela cells (mean ± SEM of n = 3 replicates; *p < 0.05).

